# Artificial light at night leads to circadian disruption in a songbird: integrated evidence from behavioural, genomic and metabolomic data

**DOI:** 10.1101/2020.12.18.423473

**Authors:** Davide M. Dominoni, Maaike de Jong, Kees van Oers, Peter O’Shaughnessy, Gavin Blackburn, Els Atema, Christa A. Mateman, Pietro B. D’Amelio, Lisa Trost, Michelle Bellingham, Jessica Clark, Marcel E. Visser, Barbara Helm

**Author notes:** Correspondence: Davide M. Dominoni.

## Abstract

Globally increasing levels of artificial light at night (ALAN) are associated with shifts in circadian rhythms of behaviour in many wild species. However, it is still unclear whether changes in behavioural timing are underlined by parallel shifts in the molecular clock, and whether such internal shifts may differ between different tissues and physiological pathways, which could highlight circadian disruption. We tackled these questions in a comprehensive study that integrated behavioural, gene expression and metabolomic analyses. We exposed captive male great tits (*Parus major*) to three ALAN intensities or to dark nights, recorded their activity rhythms and obtained mid-day and midnight samples of brain, liver, spleen and blood. ALAN advanced wake-up time, and this shift was paralleled by an advance in the expression of the clock gene *BMAL1* in all tissues, suggesting close links of brain and peripheral clock gene expression with activity rhythms. However, several metabolic and immune genes were desynchronised the shifted *BMAL1* expression, suggesting circadian disruption of behaviour and physiology. This result was reinforced by untargeted metabolomic profiling, which showed that only 9.7% of the 755 analysed metabolites followed the behavioural shift. We suggest circadian as a key mediator of the health impacts of ALAN on wild animals.

## Introduction

On our rhythmic planet, organisms have adapted to the change of day and night by evolving circadian rhythms that are highly sensitive to light [1]. The near-ubiquity of circadian rhythms across kingdoms of life suggests major fitness benefits on two grounds. Internally, the circadian system regulates temporal coordination within the body to reduce conflict and overlap between different processes. Externally, the circadian system anticipates environmental fluctuations, enabling organisms to align their behavior and physiology with nature's cycles [1,2], such as the daily alternation of light and darkness. However, globally most humans and wild organisms in their vicinity are now exposed to artificial light at night (ALAN), and thus to a rapidly altered light environment [3,4] that threatens the refined functioning of the circadian system.

In animals, rhythmicity is primarily generated on a molecular level by a transcription-translation feed-back loop (TTFL). This rhythmicity is modulated by multiple interacting systems, including neuronal, endocrine, metabolic and immune pathways [5,6][7]. The orchestration of these processes involves complex interactions between sensory input, central and peripheral clocks, and effector systems [2]. There is increasing evidence that ALAN disrupts these processes, with possible consequences ranging from compromised human health to loss of ecosystem functions [8–10]. In free-living and captive organisms, altered daily and annual activity has been widely reported, and experimental illumination has confirmed causal effects of ALAN [11,12]. Still, it is largely unclear whether the circadian system, its multiple components, and the physiological pathways it coordinates, remain synchronized with activity patterns [13– 18]. ALAN has also been shown to induce physiological changes, including in endocrine, immune and metabolic pathways [15,19,20]. These changes could be due to circadian disruption, with possible negative consequences for fitness [9,21]. Addressing these issues requires multi-level analyses that simultaneously examine effects of ALAN on rhythmic behavior and different physiological pathways [9], but these are currently lacking.

Here we aim to fill this gap by an integrated study of a bird, the great tit (*Parus major*), whose behavioral response to ALAN is well-characterized [11,22–26]. We measured day-night differences in gene transcripts in multiple tissues and in blood metabolites under a realistic range [27,28] of experimental ALAN and in dark controls, and investigated links to behavioral rhythms. The selected genes represented the circadian TTFL (Brain and Muscle ARNT-Like 1, *BMAL1*, alias *ARNTL*; cryptochrome 1, *CRY1*), a clock modulator (*casein kinase 1ε, CK1ε*) [29], and endocrine, immune and metabolic pathways putatively affected by circadian disruption (Table S1). Tissues included central pacemaker and memory sites (hypothalamus, where important avian circadian pacemaker components are located [29], and hippocampus; Fig. S1), and metabolic (liver) and immune tissues (spleen). Testes of the same birds were analyzed in a separate study [30]. In contrast to the candidate gene approach, our untargeted metabolomics approach captured both expected and novel effects of ALAN [31]. We aimed to identify whether i) hypothalamic clock gene expression was affected by ALAN, ii) potential temporal shifts in clock gene expression were consistent across tissues, iii) behavioral and clock gene rhythms were aligned, and iv) transcript and metabolite temporal shifts were consistent across physiological pathways. Any inconsistencies in temporal shifts indicate the potential for internal desynchronization, and hence, circadian disruption [9,21].

Great tits are a rewarding study system because their urbanized distribution allows to study ALAN responses also in free-living individuals, because detailed molecular and circadian information is available [32–34], and because like humans, they are diurnal [9,11,22]. We studied 34 male great tits under simulated winter daylength (LD 8.25:15.75 h) in four treatment groups, ranging from dark night controls to 5 lx (Table S2, S3), and sampled metabolites and transcripts at mid-day (3 h 30 min after lights on; i.e. 3.5 h Zeitgeber time) and midnight (7 h 15 min after lights off; i.e. 15.5 h Zeitgeber time). We chose a study design that enabled detection of rhythmicity and ALAN effects from sampling two time-points 12 h apart [35,36]. The design was enhanced firstly by tracking possible shifts in circadian rhythms by a focal clock gene, *BMAL1*, whose transcription under dark nights in songbirds peaks in the late evening [29]. Secondly, we applied ALAN levels that advance activity of captive great tits by 6 h [22] and thus, if molecular rhythms track behavior, day-night differences at all phase positions are captured.

Our specific predictions are illustrated in Figure 1, which shows expected patterns for *BMAL1*. Under dark nights (Fig. 1A, green curve), during midnight sampling (blue dots) *BMAL1* transcripts will have just passed the peak (maximum), and during mid-day (yellow dots) they will have just passed the trough (minimum). Under our hypothesis, the TTFL matches behavior, and thus, with increasing ALAN (red curves), the *BMAL1* rhythm will also advance. Hence, at midnight *BMAL1* levels will be measured progressively later than the peak, and drop, whereas mid-day levels will be measured closer to the next peak, and hence rise. When combining midnight and mid-day data (Fig. 1B), we thus expected a cross-over of detected *BMAL1* levels. Other rhythmic compounds should show similar patterns, although the point of intersection and precise change of level depends on their phase. In contrast, if the TTFL does not match the behavioral shift by ALAN, compound levels will show as two horizontal lines across ALAN, representing day and night, respectively. Levels of non-rhythmic compounds will fall on a horizonal line, representing both day and night.

**Figure 1.**
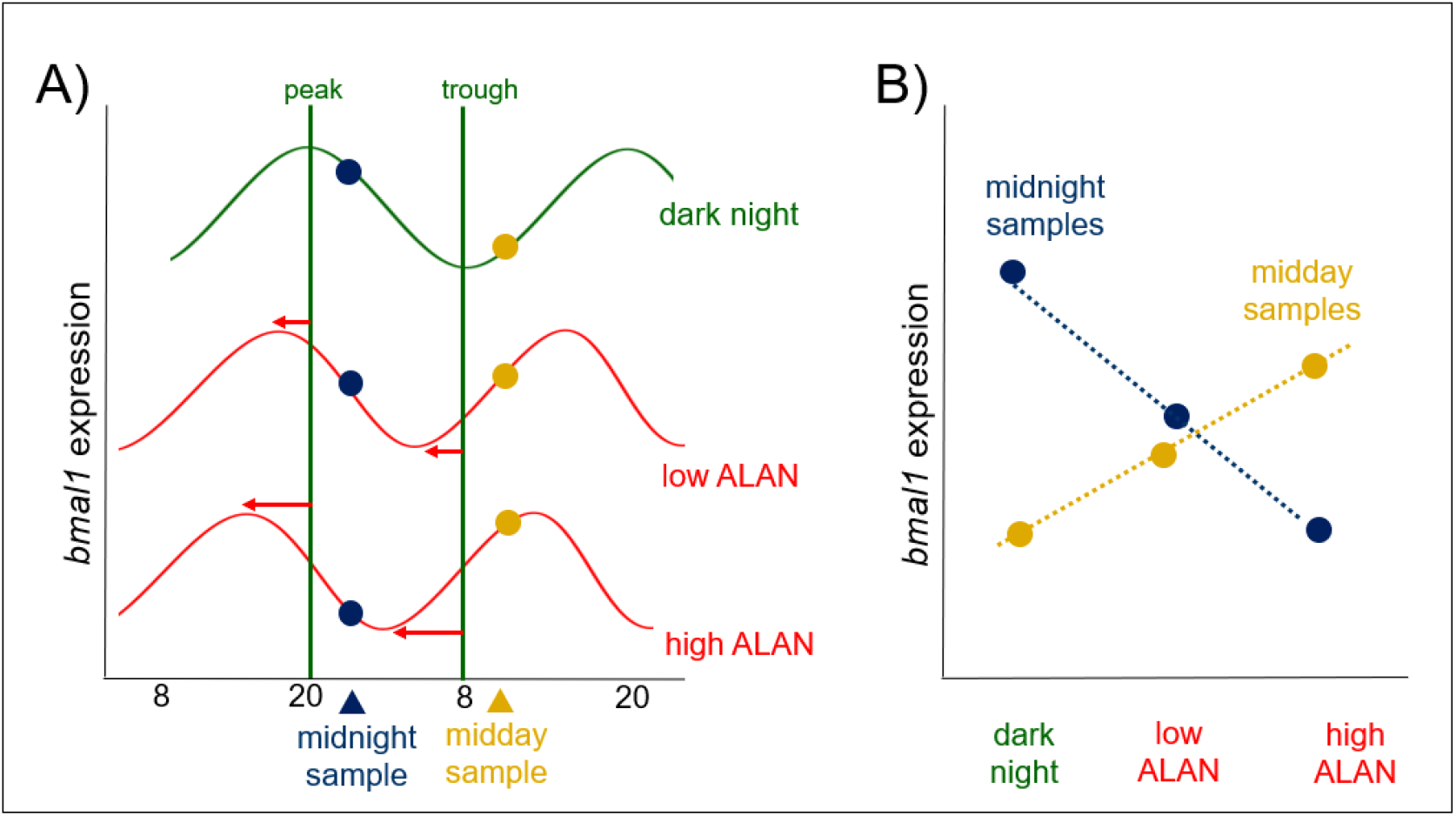
Expected clock gene rhythm advance in response to ALAN. Schematic shows ALAN effects on transcript levels of *BMAL1* measured at midnight (blue) and mid-day (yellow). (A) Rhythm of ALAN under dark night shown as green curve; if the gene's rhythm advances (red curves) with increasing ALAN, transcript levels sampled at midnight will drop, whereas those measured at mid-day will rise; horizontal arrows indicate the advance of the *BMAL1* peak. (B) The trends of transcripts with increasing ALAN therefore cross for mid-day vs. midnight sampling.

## Results

### ALAN advances circadian timing of activity and BMAL1 expression

Daily cycles of activity were strongly affected by the ALAN treatment (GAMM, p=0.001, Fig. 2A and Fig. S2; Table S4). In the 5 lx group birds were generally active 6-7 h before lights-on, whereas birds in the other two light treatments (0.5 and 1.5 lx) advanced morning activity to a much lesser extent. This advancement in the onset of morning activity led to 40% of the overall diel activity in the 5 lx group to occur during the night, compared to 11 and 14% in the 0.5 and 1.5 lx groups, and less than 1% in the control dark group. Thus, with increasing ALAN, nocturnal activity also increased (LMM, treatment p < 0.001, Fig. 2A and Table S5).

**Figure 2.**
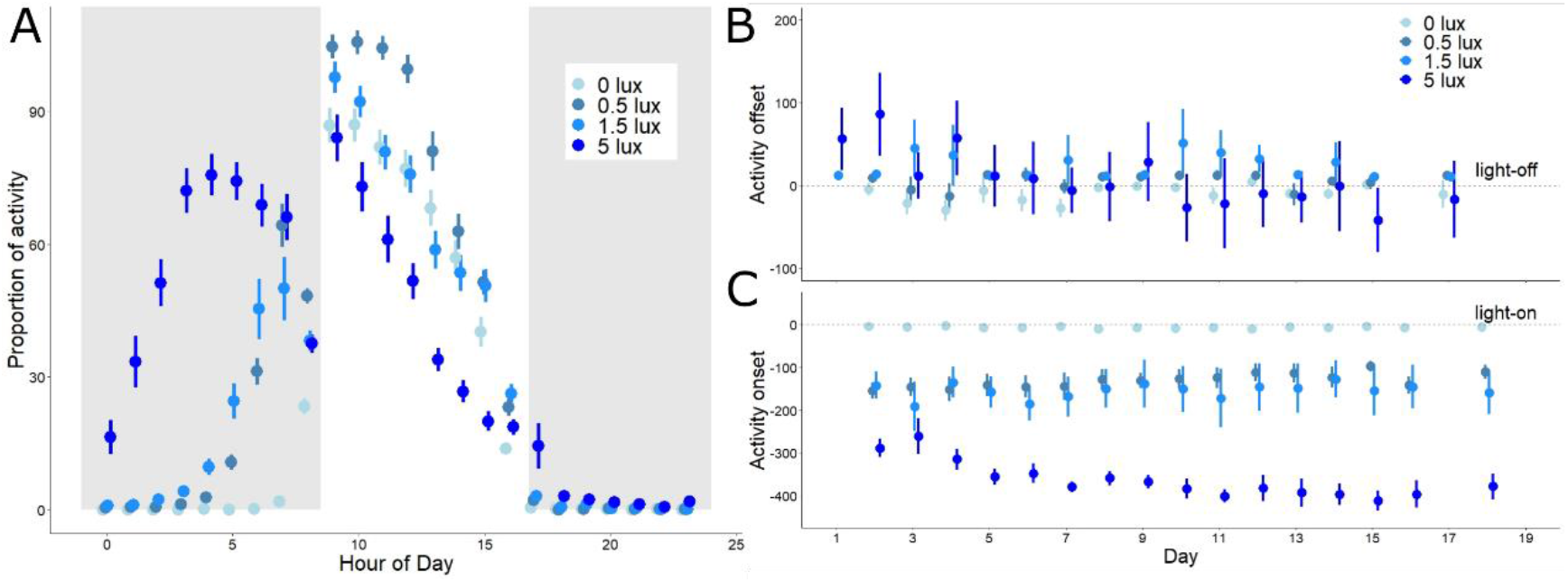
Activity timing is affected by intensity of light at night. The proportion of active 2-min intervals in each treatment group per hour of the day is shown in panel (A) (raw mean ± SEM, N = 34). Grey background indicates night-time, white background indicates daytime. On the right, we show daily treatment group data (mean ± SEM), for the timing of (A) evening offset and (B) morning onset of activity (time in min). Activity onset and offset refer to times of lights-on and lights-off, which are shown as the horizonal lines crossing zero in both panels.

Breaking down this average diel profile (Fig. 2A) by time since first exposure to ALAN (i.e., days from start of the experiment to first sampling, days 0 to 18) yields insights into how differences in activity developed, and into circadian mechanisms involved (Fig. 2B-C). Upon exposure to ALAN, the birds’ activity onset (Fig. 2C) advanced in all treatment groups. In the groups with intermediate light exposure (0.5 lx, 1.5 lx) the phase-advance occurred instantaneously and to a similar extent (155 and 142 min for the 0.5 and 1.5 lx groups respectively, P>0.1 for this pairwise comparison), but thereafter timing remained stable. The group exposed to 5 lx showed an even larger instantaneous phase advance of an average of almost five hours (mean ± SEM = 289 ± 21 min), but thereafter continued to gradually phase-advance until reaching a stable phase after 10 days (interaction treatment*day, p< 0.001, Fig. 2C, Table S2). The advance until stabilization could equally represent gradual entrainment to an early phase, or temporary free-run of activity, as suggested by periodogram analysis. Indeed, we found that in the 5 lx group, prior to stabilization, period length deviated from that of all other groups and from 24 h, reaching levels similar to those of free-running conspecifics in an earlier study [37] (mean period length 5 lx group: 23.6 h; LM; Table S6). The individual actograms (Fig. S3) further suggest that the activity rhythm in the 5 lx group may have split into an advancing morning component and a more stably entrained evening component, suggesting internal desynchronization.

Changes in the activity offset were much less pronounced (Fig. 2B). The 5 lx group showed an instantaneous phase-shift, which in contrast to morning activity delayed, rather than advanced, activity compared to the lights-off time. This initial delay was followed by a gradual advance of evening offset, similar to but smaller than that of morning onset. At the end of the experiment birds in the 5 lx group ceased their activity before lights-off, and earlier than other groups (treatment*day, p< 0.001, Fig. 2B, Table S5). This advance did not compensate for the earlier onset, as birds in the 5 lx group were more active over the whole 24h than the remaining birds (treatment*day, p= 0.01, Table S5).

### Hypothalamic BMAL1 expression at night parallels advanced activity onset

We next sought to identify whether the profound shifts in activity patterns were paralleled by corresponding shifts in the pacemaker, measured by expression of *BMAL1* in the hypothalamus. Day-night differences in transcripts of *BMAL1* inverted with increasing ALAN (Fig. S4A), as predicted above (Fig. 1). While *BMAL1* expression was higher at midnight than at mid-day for the control birds, increasing ALAN induced a reversal of this pattern, so that birds in the 5 lx group had much higher expression at mid-day than at midnight (treatment*time, p < 0.01, Table S7).

To assess whether changes in day-night *BMAL1* gene expression correlated with temporal behavioral shifts, we related *BMAL1* levels to onset of activity of an individual once it had stably shifted in response to the ALAN treatment (Fig. 2B, 2C, after 10 days). Onset was closely predicted by hypothalamic *BMAL1* expression at midnight (Gaussian LM, p<0.001, R^2^=0.71, Fig. 3A). Across ALAN levels, the earliest rising birds had the lowest midnight expression of *BMAL1*. However, the steep linear regression was largely based on differences between ALAN groups in both activity timing (Figs. 2, 3) and *BMAL1* expression (Fig. S4A). Indeed, this relationship was even stronger when we only considered the 0.5, 1.5 and 5 lx group in the analysis (Gaussian LM p<0.001, R^2^=0.85), but the association was not present for the dark control birds (Gaussian LM, P=0.87). Individual midnight *BMAL1* levels were also predictive of mean offset of activity, albeit less strongly so than for onset (Gaussian LM, p=0.006, R^2^=0.28, Fig. 3B). Conversely, mid-day *BMAL1* levels did not significantly predict variation in any of the activity traits (Gaussian LMs, p>0.1 and R^2^ <0.16 for all measures, Fig. 3C-D).

**Figure 3.**
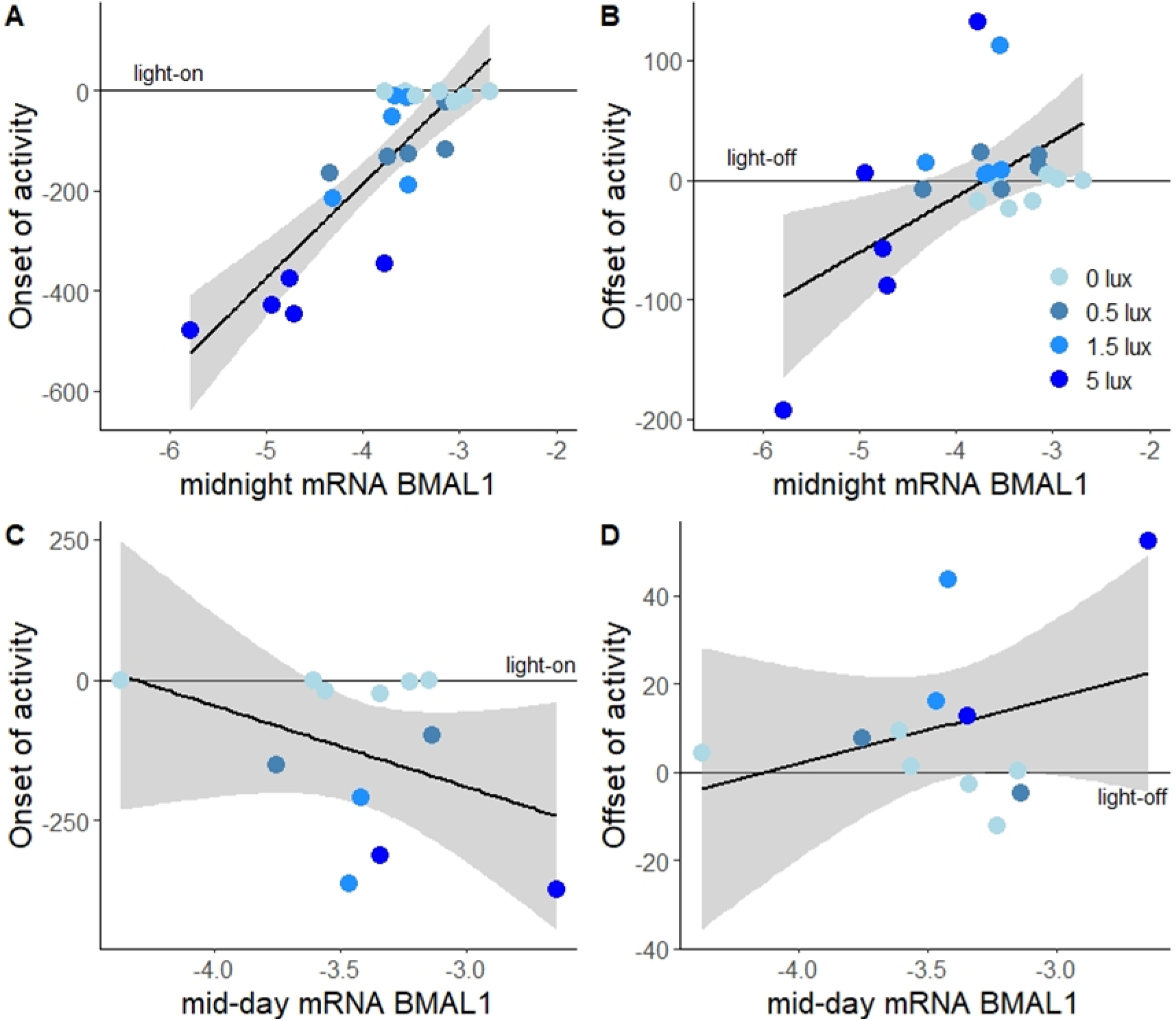
*BMAL1* expression in the hypothalamus predicts the advance of morning activity. mRNA levels of *BMAL1* at midnight correlated with the onset (A) and offset of activity (B), but mid-day levels (C, D) did not. Shown are log-transformed mRNA levels, separated by sampling time (day vs night) and ALAN treatments (blue color gradient). Points represent individual birds (total N = 34), lines and shaded areas represent model fits ± 95% confidence intervals.

### ALAN reverses day-night BMAL1 expression patterns in multiple tissues

ALAN-induced shifts in *BMAL1,* as detected in the hypothalamus, were remarkably consistent across tissues. Hippocampal *BMAL1* expression profiles resembled those in the hypothalamus (Fig. S5A) and were strongly affected by the interaction of treatment and sampling time (p<0.001, Table S8). Within individuals, mid-day and midnight transcripts in both brain tissues were closely related (LM, p<0.001, Fig. 4A, Table S9). Also liver *BMAL1* showed similar effects of ALAN on day-night expression profiles (Fig. S6A; time*treatment, p<0.001, Table S10), so that within individuals, hepatic and hypothalamic transcripts also correlated closely (LM, p<0.001, Fig. 4B, Table S9). These findings were consolidated by parallel ALAN effects on *BMAL1* expression in the spleen (Fig. S7A; time*treatment, p=0.003, Table S11), and close individual-level correlation of spleen transcripts with those in hypothalamus (LM, p=0.011, Fig. 4C) and liver (LM, p=0.001, Fig. 4D, Table S9).

**Figure 4.**
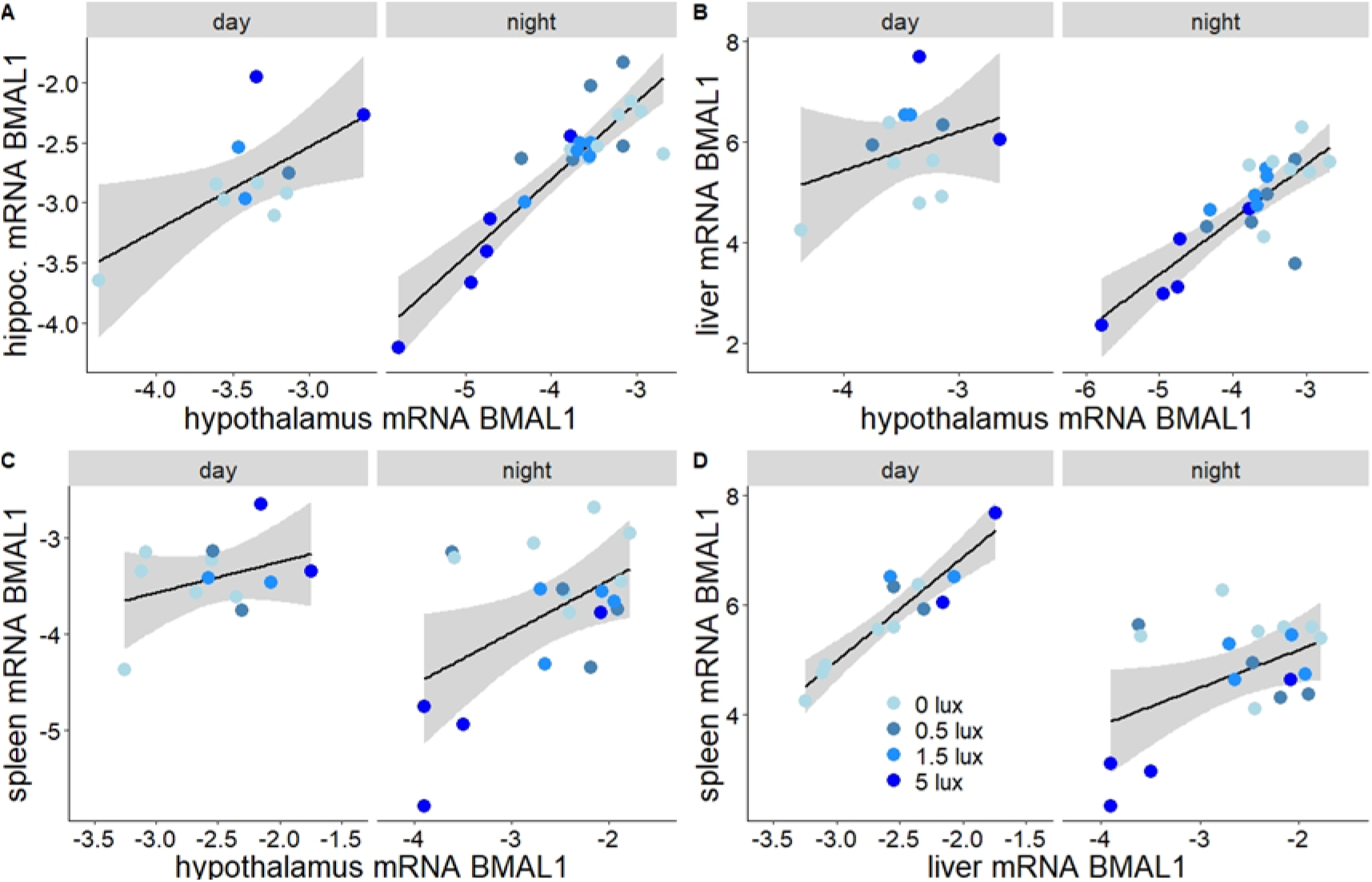
ALAN effects on *BMAL1* expression were comparable in different tissues. Correlation of expression patterns of *BMAL1* in different tissues. Shown are log-transformed mRNA levels, separated by sampling time (day vs night) and ALAN treatments (blue color gradient). Points represent individual birds (N = 34). Lines and shaded areas depict model estimated means ± 95% confidence intervals. Panels show expression levels of hypothalamic *BMAL1* levels in relation to (A) hippocampus, (B) liver and (C) spleen levels, as well as spleen in relation to liver levels (D).

### Partial disruption of expression patterns by ALAN in other genes

We next sought to assess whether the same reversal of day-night expression patterns found for *BMAL1* was paralleled in other genes analyzed in the different tissues. We found mixed evidence for this, as in most of the pathways we examined some genes shifted in concert with *BMAL1*, while others did not. This suggests that different pathways were differentially affected by ALAN.

Among clock-related genes, hypothalamic expression levels of *CK1ε*, a clock modulator, was not affected by the light treatment (p=0.71). Expression was consistently, although not significantly, higher at mid-day (p=0.09, Fig. 5H, Table S7). Similarly, the same gene was not significantly affected by sampling time or treatment in the liver. Expression of hepatic *CK1ε* increased with light intensity, albeit not significantly so (p=0.078, Fig. 5P, Table S10), and was not affected by sampling time (p=0.13, Table S10). In the liver another circadian gene, *CRY1*, showed no expression trend that aligned with that of *BMAL1* (Fig. 5O). Moreover, *CRY1* was not affected by treatment or sampling time (P>0.6 for both variables, Fig. 5O, Table S10).

**Figure 5.**
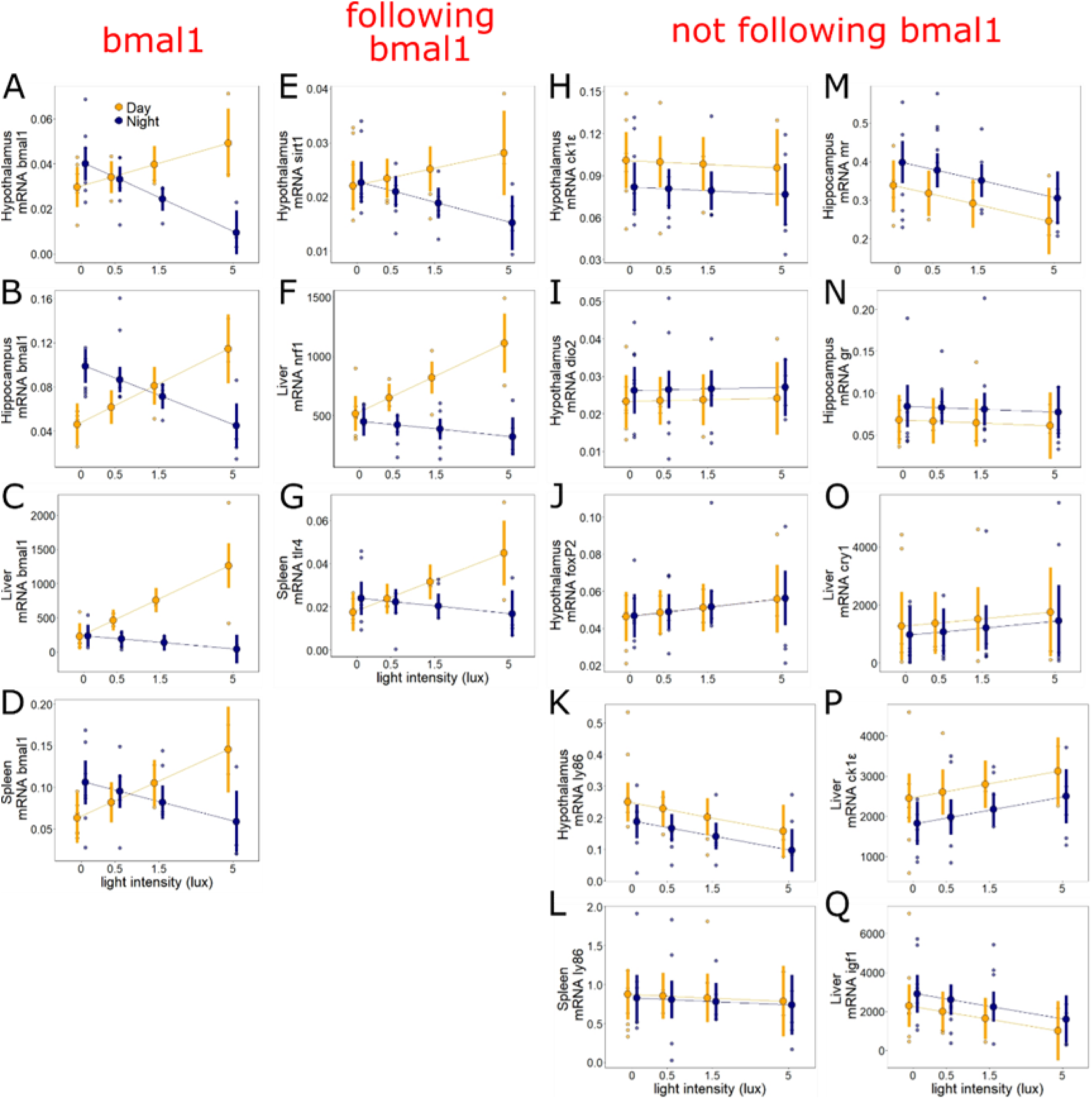
ALAN effects on gene expression are gene-specific. ALAN does not equally affect all physiological systems. ALAN effects on *BMAL1* (A)-(D) were paralleled by those on three additional genes in the hypothalamus (*SIRT1*), liver (*NRF1*) and spleen (*TLR4*) (E)-(G), but not by other genes analyzed across tissues (H)-(Q). Shown are log-transformed mRNA levels, separated by sampling time (mid-day: yellow; midnight: dark blue). Large symbols ± SEM connected by lines represent model estimates, whereas small symbols depict raw data points (N = 34 birds).

Among metabolic genes, patterns similar to those in *BMAL1* were evident in *SIRT1*, a gene which is also involved in the modulation of the circadian cycle [38][39] (Table S1). Hypothalamic *SIRT1* showed a clear change of day-night expression with increasing ALAN (Fig. 5E; treatment*time, p = 0.029, Table S7), and *SIRT1* mRNA levels were closely related to those of hypothalamic *BMAL1* (LM, p<0.001, Table S9). In the liver, the metabolic gene *NRF1* showed a similar response to ALAN as *BMAL1*, with reversed day-night expression in the 5 lx group compared to other groups (treatment*time, p<0.001, Fig. 5F, Table S10), and close correlation with *BMAL1* (LM, p<0.001). In contrast, another hepatic metabolic gene, *IGF1,* was not significantly affected by light treatment or sampling time (for both, p>0.11, Fig. 5Q, Table S10). In the hippocampus (Table S8), mid-day and midnight levels of the mineralocorticoid receptor, *MR*, decreased significantly with increasing ALAN (p=0.044, Fig. 5M). Levels were higher at night than during the day, albeit not significantly so (p=0.1). Last, the levels of the glucocorticoid receptor, *GR*, showed no significant relationship with either light treatment or sampling time (p>0.33 in both cases, Fig. 5N).

Among immune genes, ALAN affected the hypothalamic mRNA levels of *LY86*, which showed reduced levels with increasing ALAN (p=0.04, Fig. 5K, Table S7). Expression of this gene tended to be lower at midnight than mid-day, albeit not significantly so (p=0.08). However, the same gene analyzed in the spleen was not affected by either treatment or sampling time (p>0.7, Fig. 5L, Table S11). Conversely, another immune gene in the spleen, *TLR4,* showed the same pattern as *BMAL1* (Fig. 5G, time*treatment, p=0.006, Table S11).

Last, we also analyzed genes involved in photoperiod seasonal response in the avian brain. *FOXP2*, a gene that in birds is involved in learning, song development and photoperiod-dependent seasonal brain growth, showed no significant trends related to ALAN or sampling time (p>0.32 in both cases, Fig. 5J). *DIO2,* a thyroid-axis gene involved in photoperiodic reproductive activation, was also not affected by either ALAN or sampling time (p>0.45 for both variables, Fig. 5I).

### Metabolomic profiles support only a limited reversal of day-night physiology under ALAN

To explore the different impacts of ALAN on whole-body physiology, we carried out untargeted LC-MS metabolomic analysis and obtained abundance values for 5483 compounds. Out of these, 682 were annotated as known metabolites based on accurate mass and predicted retention time [40] and 73 were identified based on accurate mass measurement and matching retention time to a known standard (within 5%), for a total of 755 metabolites. We ran individual linear mixed models for all these 755 metabolites (correcting for false discovery rate at 5%), and found that 44.1% (333) differed significantly by sampling time, with higher levels at mid-day in 197, and higher levels at midnight in 136 (to see all metabolite tables: https://doi.org/10.6084/m9.figshare.12927539.v1). For 29 metabolites we found significant effects of treatment (Table S12). The direction of the treatment effect depended on the metabolite considered. In 11 metabolites, levels decreased with ALAN, while in the remaining 18 metabolites an increase was observed when compared to the dark night control group. Finally, 73 (9.7%) of the 755 metabolites showed significant interaction between treatment and sampling time (Fig. 6 and Table S13; 34 of those also differed by sampling time). As this pattern supported reversal of day-night physiology similar to that shown for activity and *BMAL1* expression, these metabolites were selected for subsequent focal analyses (hereafter named “interactive dataset”).

**Figure 6.**
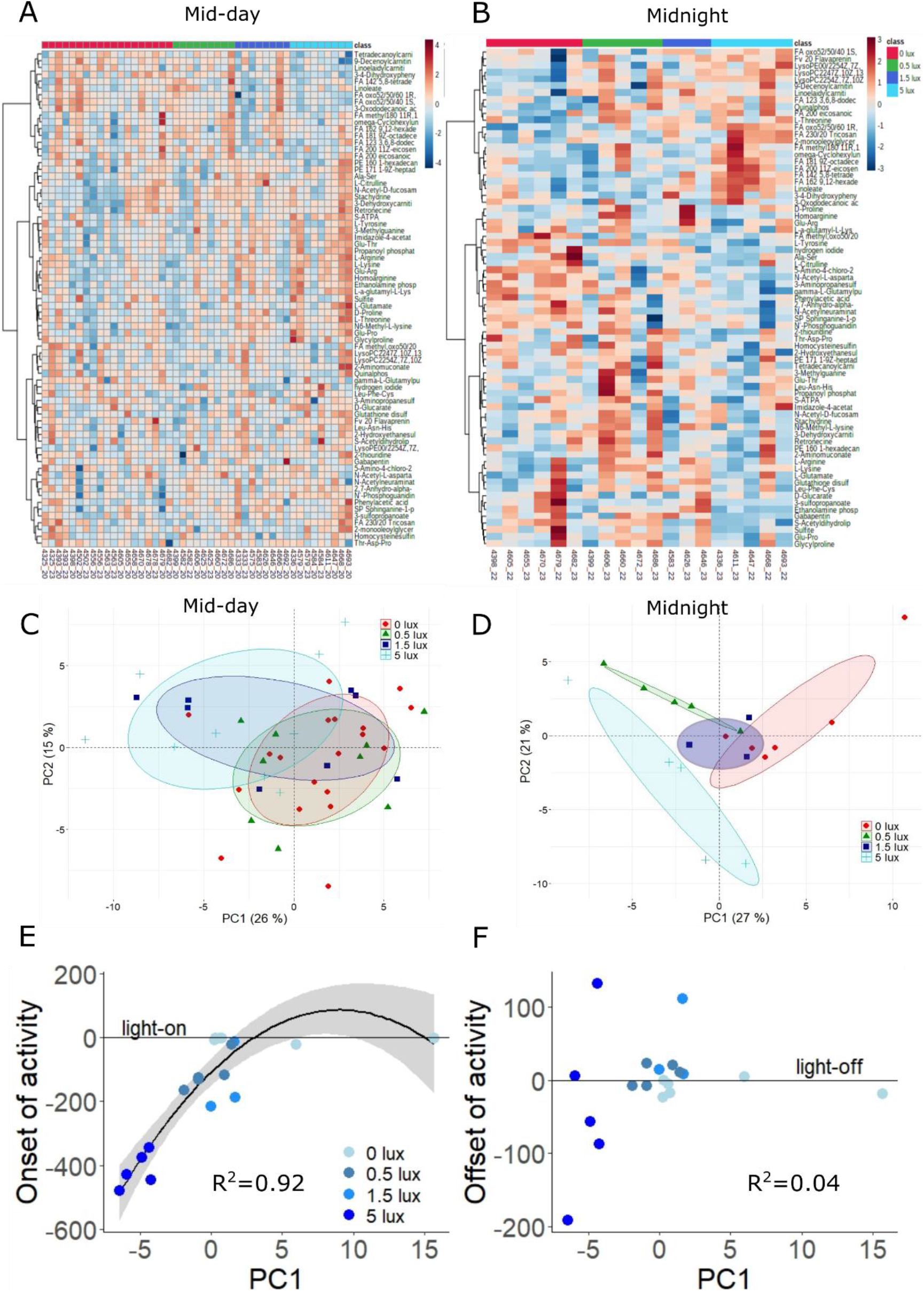
Metabolomics analysis supports ALAN-induced shifts in day-night physiology. The 73 metabolites found to be significantly affected by the interaction of treatment and sampling time (9.6% of all metabolites, interactive dataset) were dissected by means of pathway analysis and principal component analysis. Pathway analysis revealed that the Arginine Biosynthesis pathway was particularly enriched in this dataset. Heatmaps show the top-25 metabolites in the interactive dataset at either mid-day (A) or mid-night (B). Principal component analysis showed considerable overlap between ALAN groups at mid-day (C), whereas ALAN treatment effects were mostly visible at midnight, particularly for the 5 lx group (D). In all PCA plots, points represent individual samples, and ellipses contain 80% of samples in a group. The first PC of the night cluster (E) significantly predicted the onset of activity in the morning (D), but not the offset of activity in the evening (F). In (E) and (F) points represent individual birds (N = 19), and lines and shaded areas represent model fits ± 95% confidence intervals.

We dissected variation in the interactive dataset by using two principal component analyses (PCA) on the samples collected at mid-day and midnight (Fig. 6C, D). For mid-day samples, ALAN treatments overlapped considerably (Fig. 6C), although low values of PC1 (26 % of variance explained) aligned with some of the birds in the 1.5 lx and 5 lx treatments. PC1 in the mid-day dataset was heavily loaded with metabolites of Arginine biosynthesis pathway, including L-Arginine, Homoarginine and L-Glutamate, as well as other important amino acids such as L-Threonine, L-Lysine and L-Tyrosine. Conversely, the midnight samples (Fig. 6D) separated clearly between the 5 lx treatment and the remaining groups. In this midnight PCA, PC1 explained 27% of the variance and was heavily loaded with metabolites of the Glutamate and Arginine pathways, as well as with N-acetl-L-aspartate. PC2, which explained 21% of variation, was heavily loaded with fatty acids, including Linoleate (to see all factor loading tables: https://doi.org/10.6084/m9.figshare.12927536.v1). The contribution of the Arginine pathway was further confirmed by pathway analysis, conducted with Metaboanalyst [41], which indicated “Arginine biosynthesis” as a highly significant pathway in this interactive dataset (p<0.001). “Aminoacyl-tRNA metabolism” (p<0.001), “Histidine metabolism” (p=0.005), and “Alanine, Aspartate and glutamate metabolism” (p=0.026) were also indicated as significant pathways.

We finally investigated whether, just like midnight levels of *BMAL1* expression (Fig. 4), midnight principal components of metabolites correlated with individual activity timing. PC1 strongly predicted the onset of activity via a linear and quadratic relationship (n = 19, p_linear_ = 0.007, p_quadratic_=0.014, R^2^ = 0.92, Fig. 6E), but did not explain offset of activity (p=0.63, R^2^ = 0.04, Fig. 6F). PC2 was related to neither timing trait (p > 0.2).

## Discussion

Birds advanced the circadian timing of their activity as expected with increasing levels of ALAN, and in parallel the gene expression of our focal clock gene, *BMAL1,* was also advanced in the hypothalamus. Advances in *BMAL1* were consistent across tissues, indicating a shift of the circadian system in tissues implicated in timing, memory, metabolism and immune function. Furthermore, advances in nocturnal *BMAL1* potently correlated with activity onset at the individual level, consolidating close links between core clock gene expression and behavior. Responses of *BMAL1* expression were paralleled by a minority of other genes. Similarly, only 9.7% of the metabolome followed the same shift observed in *BMAL1*, indicating that most physiological pathways were desynchronized from the circadian system. The emerging picture is that birds shifted their internal clock time under ALAN, but suffered a high degree of internal desynchronization.

On a behavioral level, our findings closely match those of earlier demonstrations of advanced daily activity under ALAN in captivity for several avian species, including the great tit [15,22,24,42]. In the wild, birds also advanced daily activity under ALAN, although to a lesser extent (e.g. [14,26,43]), and often in onset but not offset [25,26,28,44,45]. Previously, behavioral shifts were interpreted as not involving the circadian clock [24]. In an experiment also on the great tit, Spoelstra and colleagues [24] exposed birds to dark nights and then to ALAN as in our study. Subsequently, birds were released to constant low-levels of dim light (0.5 lx), where they free-ran. The study found that the birds free-ran from the timing they had shown under initial dark nights, rather than from their advanced timing under ALAN. Thus, the authors concluded that the behavioral response to ALAN was due to masking, while the internal clock remained unchanged [24]. Our molecular data suggest a different conclusion, namely that within three weeks of ALAN exposure, internal time had phase-advanced in concert with behavior. These discrepancies are difficult to interpret because inferences of the studies are based on different criteria (molecular vs. behavioral) and different experimental phases (during ALAN vs. during ensuing free-run), but it is clear that additional experimental data are needed.

Our transcriptional findings of ALAN-altered rhythmicity gain support from a comparison of clock gene expression in Tree sparrows (*Passer montanus*) from an illuminated urban and dark non-urban habitat [46]. Sampled within a day after being brought into captivity, urban birds showed clear advances in the circadian system, including, as in our birds, in hypothalamic *BMAL1*. Other experimental studies have also confirmed effects of ALAN on avian rhythms in brain and other tissues [16,17]. In our study, only some of the investigated regulatory genes aligned with the ALAN-dependent advances of rhythms in behavior and *BMAL1*. The genes from metabolic pathways that have close molecular links to the TTFL, *SIRT1* and *NRF1*, mirrored ALAN-dependent changes in *BMAL1*. However, regulatory genes of immune pathways responded inconsistently, whereby *TLR4* aligned with *BMAL1* whereas *LY86* did not. The learning gene, *FOXP2* and the thyroid-activating gene *DIO2* did not mirror the changes in *BMAL1*, nor did the endocrine genes (*MR, GR, IGF1*). Conversely, in a complementary study on these same birds, we observed that ALAN exposure, which also activated the reproductive system, shifted the day-night expression patterns of corticoid receptors [30].

Other experimental studies have confirmed that effects of ALAN on avian rhythms in brain and other tissues differed between genes and pathways. For example, a study on Zebra finches (*Taeniopygia guttata*) reported ALAN-induced changes in rhythmic expression of hypothalamic *CRY1* but not *BMAL1* [16]. This differs from our findings, where advances in *BMAL1* were not paralleled by *CRY1* [17], and from findings that *BMAL1* and *CRY1*, but not another TTFL gene, *CLOCK*, advanced in an urban bird [47]. Divergent responses between clock genes might participate in circadian disruption, and could underlie discrepant behavioral responses, such as differences between activity onset and offset observed in our study, and in wild great tits [25,26,44] and other avian species [28,45]. In our study in the 5 lx group, we also observed splitting of rhythms, which has previously been linked to reproductive activation [48], a known side-effect of ALAN [13].

Our metabolomic data corroborated our main findings on gene expression. Of the 755 identified metabolites, nearly 50% (333) differed between mid-day and mid-night levels. However, less than 10 % showed changes in rhythm under ALAN (Fig. 7). These findings confirm that some, but not all featured pathways aligned with shifts in behavior and *BMAL1*. Our findings from captive wild birds under ALAN match those from human studies. To identify the mechanisms by which circadian disruption drives metabolic disorders and other pathologies, these studies severely disrupted the circadian system by sleep deprivation and shift-work protocols [31,49,50]. The reported changes in gene expression and metabolite levels were similar to those of our birds under ALAN, including highly responsive pathways and compounds, in particular Arginine [50], an amino acid strongly linked to circadian rhythms and innate immune responses [51]. Glutamate production from arginine is well known [52], and changes in these two metabolites may be due to changes in energy requirements at the different light intensities. N-acetyl-aspartate, a metabolite involved in energy production from glutamate [53], was also observed to follow changes in behavior and *BMAL1*. Both glutamate and arginine have a variety of biochemical roles [54,55], so further work would be required to determine which of these functions, if any, are associated to the behavioral and gene expression changes we observed. While preliminary, this data shows the potential of metabolomic techniques for furthering this area of research.

**Figure 7.**
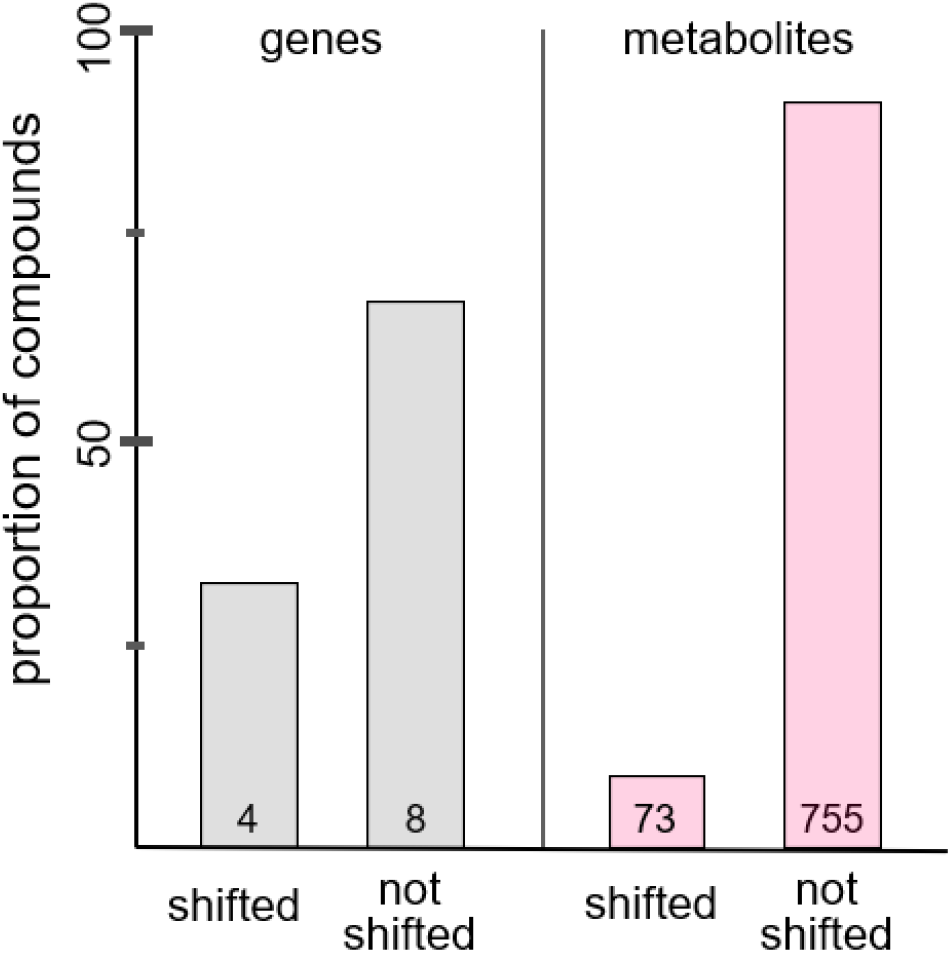
Proportion of shifts in day-night pattern in response to ALAN. Shown are proportions of genes (grey) and metabolites (red) whose levels were, or were not, significantly impacted by the interaction of sampling time and ALAN level.

Despite our sampling design of only two time-points and low sample sizes, we derived descriptors of internal time (*BMAL1* expression; metabolomics PC1 of interactive dataset) whose midnight levels had high predictive power of activity timing. Thereby, we have shown that internal time can be captured in birds by a single sample of blood or tissue, a frontline ambition of biomedical research [35,36]. Our predictive power was limited to treatment groups and within-ALAN individuals, whereas birds kept under dark nights were highly synchronized to the sudden switch of lights-on.

For wild animals, our study adds to emerging evidence of detrimental effects of ALAN on physiological pathways [9,10,21]. For example, under ALAN molecular markers for sleep deprivation were elevated, hypothalamic expression of genes such as *TLR4* was altered [16], neuronal features in the brain were changed, and cognitive processes and mental health-like states were impaired [16,20,56,57]. Altered hepatic expression of several metabolic genes further suggested negative effects on gluconeogenesis and cholesterol biosynthesis [15]. Consequences of ALAN-induced changes in immune function include increased host competence for infectious disease [58], indicating how effects on individuals may cascade to ecological or epidemiological scales.

Addressing effects of ALAN is therefore urgent [10,59]. Our data contribute to the rising evidence for dose-dependent responses of behavior and physiology [22,30,60], which might allow mitigating against ALAN impacts on wildlife by reducing light intensity [61]. Importantly, we detected substantial effects even at light intensities (0.5 lx) that are typically far exceeded by street illumination, and to which animals are exposed to in the wild [27,28]. These findings transfer to other organisms including plants, insects, and mammals including humans [12,62–65] and call for limits to the ever faster global increase in light pollution [3].

## Methods

### Data availability

The full details of our methods are presented in the *Supporting* Information document. Raw data, created datasets and R scripts are available via Figshare: (https://figshare.com/projects/Artificial_light_at_night_shifts_the_circadian_system_but_still_leads_to_physiological_disruption_in_a_wild_bird/88841).

### Animals and experimental design

We studied 34 hand-raised, adult male great tits that were kept in individual cages (90 × 50 × 40 cm) under simulated natural daylength and ambient temperature of 10 to 14 °C with *ad libitum* access to food and water, as described in [30].

The experiment started on February 1^st^, 2014, when daylength was fixed at 8 h 15 min light and 15 h 45 min darkness. During the day, all birds were exposed to full spectrum daylight by high frequency fluorescent lights emitting ~1000 lx at perch level (Activa 172, Philips, Eindhoven, the Netherlands). During the night, birds were assigned to four treatment groups exposed to nocturnal light intensity of 0 lx (n= 13), 0.5 lx (n = 7), 1.5 lx (n = 7), or 5 lx (n = 7). In composing these groups, we prioritized assigning birds to the dark night group to obtain reliable benchmark data on day-night differences in gene expression. Lights were provided by warm white LED light (Philips, Eindhoven, The Netherlands; for details on the spectral composition of lights, see [22]).

On Feb 20^th^ an initial blood sample (~200 μl) was collected from all birds at mid-day for metabolomic profiling. On Feb 22^nd^ birds were randomly assigned to mid-day or midnight groups for culling to collect tissues for morphological and molecular analyses. The mid-day group was culled on Feb 22^nd^, whereas culling of the midnight group was divided over two subsequent nights (Feb 22^nd^: 12 birds; Feb 23^rd^: 10 birds). Blood was again collected for metabolomic profiling.

All experimental procedures were carried out under license NIOO 13.11 of the Animal Experimentation Committee (DEC) of the Royal Netherlands Academy of Arts and Sciences.

### Locomotor activity

Daily activity patterns of each individual bird were measured continuously using micro-switches recorded by a computer, as described in de Jong et al [22]. See Supporting Information for more details.

### Gene expression analyses

After culling, organs were extracted, snap-frozen on dry ice, and stored at −80 °C within 10 min of capture.

Brain tissue was cut on a cryostat at – 20 °C. We cut sagittal sections throughout the brain (Fig. S1). The hypothalamus and hippocampus were located by the use of the Zebrafinch atlas ZEBrA (Oregon Health & Science University, Portland, OR, USA; http://www.zebrafinchatlas.org) and isolated from the frozen brain sections either by surgical punches for the hypothalamus (Harris Uni-Core, 3.0 mm), or by scraping the relevant tissue with forceps, for the hippocampus. For the hypothalamus, the edge of the circular punch was positioned adjacent to the midline and ventral edge of the section, just above the optic chiasm, following the procedure of [66]. Hypothalamic and hippocampal tissue was then immediately added to separate 1.5ml buffer tubes provided by the Qiagen RNeasy micro extraction kit (see below), homogenized and stored at −80 °C until extraction.

Whole spleens were homogenized with a ryboliser and added to 1.5 ml RNeasy micro buffer and stored at −80 °C. For livers, we cut 0.5 g of tissue from each individual liver, homogenized it and added it to 1.5 ml RNeasy micro buffer and stored it at −80 °C. RNA was extracted using the RNeasy micro extraction kit and reverse transcribed it to generate cDNA using a standard kit following the manufacturer’s instructions (Superscript III, Invitrogen).

We selected exemplary genes known to be involved in circadian timing, seasonal timing, and in metabolic, immune and endocrine function (Table S1). We analyzed the core clock gene *BMAL1* in all tissues as our primary clock indicator because of the timing of its expression and because of its role as central hub for inter-linking molecular pathways [7]. We also studied a second core clock gene, *CRY1*, in a single tissue, and a clock modulator, *CK1ε*, in two tissues. In the hypothalamus, we also studied two genes involved in seasonal changes (*DIO2, FOXP2*), and one metabolic and ageing gene (*SIRT1*). The second metabolic gene, *NRF1*, was studied in the liver. Two immune genes represented different pathways (*LY86, TLR4*). Finally, we studied endocrine genes involved in stress signaling in the Hippocampus (*NR3C1 (alias GR), NR3C2 (alias MR)*) and in tissue homeostasis (*IGF1*), as well as reference genes (for full details see Table S1). Primers were built based on the great tit reference genome build 1.1 (https://www.ncbi.nlm.nih.gov/assembly/GCF_001522545.2) [33] and annotation release 101 (https://www.ncbi.nlm.nih.gov/genome/annotation_euk/Parus_major/101/). Primer design was conducted with Geneious version 10.0.2 [67].

Amplification efficiency of each primer pair was determined through quantitative real-time polymerase chain reaction (RT-qPCR). RT-qPCR was performed on duplicate samples by a 5-point standard curve. We used reference gene levels to correct for variation in PCR efficiency between samples. Reference gene expression stability was calculated using the application geNorm [68], from which we identified the best pair of reference genes for each tissue. Absolute amounts of cDNA were calculated by conversion of the Ct values (C×E^−Ct^, with C=10^10^ and E=2) [69]. The absolute amounts of the candidate genes were then normalized by division by the geometric mean of the absolute amounts of the reference genes. This step yielded relative mRNA expression levels of the candidate genes. For more details, see the Supporting Information document.

### Metabolomics analysis

See Supporting information for initial sample preparation and for additional details. All samples were analyzed on a Thermo Scientific QExactive Orbitrap mass spectrometer running in positive/negative switching mode. Mass spectrometry data were processed using a combination of XCMS 3.2.0 and MZMatch.R 1.0–4 [70]. Unique signals were extracted using the centwave algorithm [71] and matched across biological replicates based on mass to charge ratio and retention time. The final peak set was converted to text for use with IDEOM v18 [72], and filtered on the basis of signal to noise score, minimum intensity and minimum detections, resulting in a final dataset of 755 metabolites.

### Statistical analysis

All statistical analyses were conducted in R, version 3.63 [73]. In all models we included treatment as log-transformed light intensity (adding a constant to avoid zero). Details of all statistical analyses can be seen in the Supporting Information document.

To analyze locomotor activity data (i.e. perch-hopping), we first divided the time series of activity into an unstable phase and stable phase (see Supporting information). We used the data in the unstable phase to quantify circadian period length (tau) for each bird, then tested treatment effects using a gaussian linear model (LM). The data in the stable phase were used to test for variation in the proportion of time spent active every hour depending on treatment, using a generalized additive mixed model (GAMM). Finally, we tested for variation in onset time, offset time, nocturnal activity and total daily activity using separate linear mixed models (LMMs).

To examine variation in relative transcript levels, we ran LMs including ALAN treatment, sampling time (two-level factor, day and night), and their interaction as explanatory variables, and mRNA expression levels of the different genes in the different tissues as response variables. Similar models were used to test for relationships in mRNA levels between the same gene in different tissues, or different genes in the same tissue.

To test for variation in the levels of the individual metabolites identified by the LC-MS, we used all data, including the replicated mid-day samples (total n = 64). We ran independent LMMs for each metabolite, with metabolite levels as response variable (log transformed and normalized), and treatment, time of day and their interaction as explanatory variables. Moreover, we ran two principal component analyses using only the 73 metabolites found to be significantly affected by the treatment*time interaction in the LMMs described above. The two PCAs were run separately for the individual samples collected at mid-day or midnight. We then used the first two principal components (PC1 and PC2) of the midnight based PCA as explanatory variables in two LMs with onset and offset of activity as response variables, respectively.

## Acknowledgments

This work was supported by a Wellcome Trust grant to B.H and D.M.D (097821/Z/11/Z), a Marie-Curie Career Integration Grant to B.H. (ECCIG (618578) Wildclocks), a NERC Highlight Topics grant to D.M.D. (NE/S005773/1) and the Dutch Technology Foundation (STW). We thank Kamiel Spoelstra, Takashi Yoshimura and Bill Schwartz for fruitful discussions on the results of this study.

## Author Contributions

DMD, MdJ, MEV and BH designed the study. DMD, MdJ, PBD, LT and BH collected the data and samples. DMD, KvO, POS, EA, CAM, MB, JC performed the gene expression assays. GB performed the metabolomics analyses. DMD conducted the statistical analyses. DMD and BH wrote the paper.

## Competing Interest Statement

We declare no competing interests.

## Notes

https://figshare.com/projects/Artificial_light_at_night_shifts_the_circadian_system_but_still_leads_to_physiological_disruption_in_a_wild_bird/88841

### Competing Interest Statement

The authors have declared no competing interest.

### Summary of Updates

Title and abstract. Supplementary materials have also been attached.

