## Supplementary methods, tables and figures for "Artificial light at night leads to circadian disruption in a songbird: integrated evidence from behavioural, genomic and metabolomic data"

All datasets and R scripts to reproduce the results of this study are available in Figshare: (https://figshare.com/projects/Artificial_light_at_night_shifts_the_circadian_system_but_still_leads_to_physiological_disruption_in_a_wild_bird/88841).

### Supplementary material and methods

*Animals and experimental design*

We conducted the experiment between February 1 and February 23, 2014, as described elsewhere for these same birds [1]. We used 34 adult male great tits that had been used in a previous experiment aimed at assessing the impact of different levels of light intensity at night on daily activity and physiology [2]. All birds had been hand-raised and housed at the Netherlands Institute of Ecology (NIOO KNAW), Wageningen, The Netherlands, in indoor-facilities in individual cages (90 × 50 × 40 cm). All birds were between 1 and 4 years of age (hatched in 2012 or before), but mean age did not differ significantly between treatment (P = 0.576). Temperature was maintained between 10 and 14 °C, and did not vary between day- and night-time. Birds had access to food and water ad libitum. We used dividers between the cages, so that birds could only hear but not see each other, and light from one cage did not influence the light environment in adjacent cages.

During the ALAN experiment, birds were kept under fixed natural day-length of 8 hr 15 min light and 15 hr 45 min darkness. Each cage had two separate light sources for day- and night-time illumination. During the day, all birds were exposed to full spectrum daylight by high frequency fluorescent lights emitting ~1000 lux at perch level (Activa 172, Philips, Eindhoven, the Netherlands). For night-time, birds were assigned to different treatment groups that varied in the level of light intensity used (warm white LED light; Philips, Eindhoven, The Netherlands). The spectral composition of this light is shown in Supporting Information Figure S1 of [2], based on an earlier experiment with these birds. In this earlier experiment, the birds were exposed to five levels of ALAN for one month between December 10, 2013 and January 10, 2014 and otherwise kept under dark nights. The experimental setup we used here differed as we used four, and not five experimental levels of ALAN, of which one now was a dark control (0.00, 0.5, 1.5, and 5 lux). The birds in the dark control were derived from the two earlier treatment groups with the lowest light intensity (0.05 and 0.15 lux, respectively), while birds in all other treatments were kept in the same treatment that they were exposed to in the previous experiment. Thus, from the start of our present experiment on February 1, 2014, the birds were exposed for the entire night to either one out of three nocturnal light intensities measured at perch level in the cages: 0.5 lux (n = 7), 1.5 lux (n = 7), or 5 lux (n = 7), or to dark control conditions (n = 13) (Table S2).

The four treatment groups were assigned to one of seven blocks of cages arranged within two experimental rooms. Each block contained all treatment groups, distributed using a Latin Squares design. The birds were kept under these conditions for 3 weeks until culling to collect tissues for morphological, metabolomic and genetic analyses (see more details on this terminal sampling below). On Feb 20$^{th}$ we collected a blood sample (~200 µl) from all birds for metabolomic profiling. The randomly assigned mid-day sampling group was culled on Feb 22$^{nd}$, and the midnight group during the two subsequent nights (Feb 22$^{nd}$: 10 birds; Feb 23$^{rd}$: 12 birds). All experimental procedures were carried out under license NIOO 13.11 of the Animal Experimentation Committee (DEC) of the Royal Netherlands Academy of Arts and Sciences.

*Locomotor activity*

A standard wooden perch and a perch with a micro-switch were fitted into every cage before the start of the experiments. The micro-switch detected perch-hopping and logged the frequency onto a computer. A signal for on (bird on perch) and off (bird not on perch) was recorded every 0.1 s and stored in files as 30 s intervals by software developed by T&M Automation (Leidschendam, The Netherlands). An activity level of either one or zero was obtained for every two minutes, in which a bird was considered active if the micro-switch was triggered once or more times.

In total, we examined four different aspects of activity for each bird over a 24-h period with the program Chronoshop 1.1 (by K. Spoelstra). Activity onset (the first time point at which activity is higher than the average) and activity offset (the final time point which activity is higher than the average) were reported relative to when the daylight was switched on and off, respectively. Total activity was defined as the total active minutes within a 24-h cycle (from midnight to midnight), while nocturnal activity was defined as the total number of active minutes during the relative night (lights off until lights on).

*Tissue preparation*

We culled birds under isoflurane anesthesia (Forene, Abbott, Hoofddorp, the Netherlands) at mid-day (±2 hr) on February 22, 2014 or midnight (±2 hr) on February 22 and 23, 2014. Organs were extracted, snap-frozen on dry ice, and stored at −80 ºC within 10 min of capture. The final sample size is shown in Table S2.

The whole brain was cut sagittally on a cryostat at – 20 ºC, alternating three sections of 40 µm with one section of 60 µm. The 40 µm sections were used to collect tissue for gene expression analysis, thus after being cut they were temporarily stored again at −80 ºC until RNA extraction. The 60-µm sections were immediately Nissl-stained and used as reference, to verify histologically that we collected tissue from the regions of interest in the 40 µm sections. From the appearance of the cerebellum in the slides, we collected 120 slices until the disappearance of the cerebellum at the opposite side, using a total of 90 slices for RNA extraction and 30 slices as reference. The hypothalamus and hippocampus in the Nissl-stained series were identified by referencing the Zebrafinch atlas ZEBrA (Oregon Health & Science University, Portland, OR, USA; http://www.zebrafinchatlas.org). To isolate the tissue, for the hypothalamus we sampled one 3 mm of diameter circular tissue punch (Harris Uni-core, Electron Microscopy Sciences, cat#69036) from each section. We used the optic chiasma, medially, the dorsal supraoptic decussation (rostrally, when visible), and the optic tract, laterally, as a reference and punched the medial area immediately dorso-caudal to the optic chiasma (Fig. S1). The collected areas corresponded roughly to the medial basal suprachiasmatic hypothalamus region which is comprised of several nuclei and areas including the suprachiasmatic nuclei (SCN). For the hippocampus, we used forceps to remove the tissue from the medial (enlarged) part of the hippocampus above the lateral ventricle up to the edge of the brain (Fig. S1). Hypothalamic and hippocampal tissues were then immediately added to separate 1.5ml buffer tubes provided by the Qiagen RNeasy micro extraction kit (see below), homogenized and stored at -80 °C until extraction.

Whole spleens were homogenized with a ryboliser and added to 1.5 ml RNeasy micro buffer and stored at -80 ºC. For livers, we cut 0.5 g of tissue from each individual liver, homogenized it and added it to 1.5 ml RNeasy micro buffer and stored them at -80 ºC.

*RNA isolation and cDNA synthesis*

RNA isolation and cDNA syntheses was conducted in Glasgow. RNA was extracted using the RNeasy micro extraction kit (Qiagen) following the manufacturer's protocol. RNA quality and quantity were evaluated using a NanoDrop 2000 spectrophotometer (ThermoFisher Scientific). RNA yield was used to adjust the concentration for cDNA synthesis. The working RNA concentration was 25 ng/µl. For each tissue sample, we used 6µl of RNA and reverse transcribed it to generate cDNA using a standard kit following the manufacturer's instructions (Superscript III, Invitrogen). We tested serially diluted cDNA samples for each gene of interest to determine an optimal dilution.

*Primer design*

We made a list of genes known to be involved in circadian and seasonal timing, as well as in metabolism and immune function (Table S1). Similarly, we made a list of reference "housekeeping" genes to allow normalization of the gene expression. Primers were built based on the great tit reference genome build 1.1 (https://www.ncbi.nlm.nih.gov/assembly/GCF_001522545.2 ) [3] and annotation release 101 (https://www.ncbi.nlm.nih.gov/genome/annotation_euk/Parus_major/101/). Primer design was conducted with Geneious version 10.0.2 [4]. Primers were checked against the great tit reference genome using a BLAST search to confirm that primers were specific for the intended target genes. In order to avoid genomic DNA (gDNA) amplification, every primer pair was designed to span an intron of more than 1000 base pairs. Selected primer pairs for the final candidate genes are listed in Table S1.

*RT-qPCR*

Amplification efficiency of each primer pair was determined through quantitative real-time polymerase chain reaction (RT-qPCR). We first analyzed liver samples in Glasgow, measuring fluorescence with a MX3000 cycler 96-well plates (Stratagene). By the time we could analyze brain and spleen samples, logistic issues prevented us to run these assays in Glasgow. Thus, we proceeded with analyses at the NIOO in Wageningen (see below for validation), where fluorescence was measured with the CFX Connect Real-Time PCR Detection System (Bio-Rad Laboratories). In both cases, RT-qPCR was performed by a 5-point standard curve based on a 5-dilution series (1:10, 1:20, 1:40, 1:80 and 1:160) of cDNA samples. We included duplicated samples (10 µl) for one transcript on each plate, balancing time points and treatments, using the SYBR Green method (PowerUp SYBR Green Master Mix, ThermoFisher Scientific). At the end of the amplification phase, a melting curve analysis was carried out on the products formed. In each plate, we also included duplicate negative control wells (with RNA instead of cDNA). None of the primer pairs amplified gDNA and the efficiency of the qPCR reactions was always between 95% and 103%.

The PCR efficiency and fractional cycle threshold number obtained in Glasgow and Wageningen were used for gene quantification. We used reference gene levels to correct for variation in PCR efficiency and RNA quality between samples. From our list of starting reference genes, we selected two per tissue to correct candidate (target) gene levels. Reference gene expression stability was calculated using the application geNorm [5], from which we identified the best pair of reference genes. Absolute amounts of cDNA were calculated by conversion of the Ct values ($C \times E^{-Ct}$, with $C=10^{10}$ and $E=2$) [6]. The absolute amounts of the candidate genes were normalized by division by a normalization factor, calculated by taking the geometric mean from the absolute amounts of the reference genes. We thereby obtained relative mRNA transcript levels of the candidate genes.

*Validation of qPCR data obtained in Glasgow and Wageningen*

As quantification of qPCR products might be influenced by the machine used for the analyses, we decided to validate the data produced at the two different laboratories by analysing liver *bmal1* levels in Wageningen, too. Correlation between Glasgow and Wageningen liver data was highly significant (Spearman rho: 0.74, $p<0.001$).

*Metabolomics*

The 68 plasma samples (34 individuals x 2 time points = 68) were first prepared by the following protocol adjusted for the amount of plasma available (minimally 26 µl): 100 µl of plasma were mixed with chloroform and methanol in a 1:3:1 ratio (chloroform : methanol : sample) on a cooled shaker for one hour and then centrifuged for 3 mins at 13,000g at 4 °C. The resulting supernatant was stored at -80 °C until LC/MC analysis. All samples were analyzed on a Thermo Scientific QExactive Orbitrap mass spectrometer running in positive/negative switching mode. This was connected to a Dionex UltiMate 3000 RSLC system (Thermo Fisher Scientific, Hemel Hempstead, United Kingdom) using a ZIC-pHILIC column (150 mm × 4.6 mm, 5 µm column, Merck Sequant, Gillingham, UK). The column was maintained at 30 °C and samples were eluted with a linear gradient (20 mM ammonium carbonate in water, A and acetonitrile, B) over 46 min at a flow rate of 0.3 mL/min as follows 0 min 20% A, 30 min 80% A, 31 min 92% A, 36 min 92% A, 37 min 20% A, 46 min 20% A. The injection volume was 10 µL and samples were maintained at 5 °C prior to injection.

Mass spectrometry data were processed using a combination of XCMS 3.2.0 and MZMatch.R 1.0–4 [7]. Briefly, data were converted from Thermo proprietary raw files to the open format mzXML. Unique signals were extracted using the centwave [8] algorithm and matched across biological replicates based on mass to charge ratio and retention time. These grouped peaks were then filtered based on relative standard deviation and combined into a single file. The combined sets were then filtered on signal to noise score, minimum intensity and minimum detections, leading to the exclusion of four bird samples (final n = 64). The final peak set was then gap-filled and converted to text for use with IDEOM v18 [9]. IDEOM contained peak data from 5483 compounds. From this dataset, we removed all compounds that IDEOM defined having a confidence lower than 4 (out of 10). The majority of these were fragments that did not match to any known standard mass and retention time. Moreover, some metabolites were present twice as two different polarities, in which case only the polarity returning the highest signal was maintained, while the other was removed. Last, metabolites with 0 abundance in more than 10 % of the individuals were also removed. The final dataset contained 755 metabolites.

*Statistical analysis*

All statistical analyses were conducted in R, version 3.63 [10]. In all models we included treatment as log-transformed light intensity (adding a constant = 1 to avoid zero returns).

To analyze locomotor activity data (i.e. perch-hopping), we first divided the time series of activity into a first phase, before all birds had stabilized their activity timing (defining the day of stabilization as the first day when the mean onset did not differ for more than 1 SEM to the previous and following day), and a stabilized phase thereafter. We first estimated the free-running period of birds during the first 10 days, by calculating the circadian period length using the Lomb-Scargle periodogram analysis implemented in the software Chronoshop (credits to Kamiel Spoelstra). We tested for differences in circadian period length between groups using a Gaussian LM with treatment as explanatory variable.

Second, using only the stabilized activity data after day 10, we ran a generalized additive mixed model (GAMM) to test for variation in the proportion of time spent active every hour. Bird ID was included as random factor. We included hour of day, in interaction with treatment, as smoothed terms. The GAMM was run using the function gamm in the package *mgcv* [11].

Third, we tested for variation in onset time, offset time, nocturnal activity and total daily activity using separate linear mixed models (LMMs) with ID as random effect, and treatment, day of the experiment and their interaction, as well as the quadratic effect of day, as explanatory variables. LMMs were run using the function lmer in the package *lme* [12].

To examine variation in relative transcript levels, we ran linear models (LMs) including ALAN treatment, sampling time (two-level factor, day and night), and their interaction as explanatory variables, and mRNA expression levels of the different genes in the different tissues as response variables. We used linear models also to test for individual-level relationships between mRNA levels of the same gene in different tissues, or for the relationship between mRNA levels of different genes withing the same tissue. Moreover, we also used linear models to relate midnight hypothalamic mRNA levels of BMAL1 (explanatory variable) to the mean time of activity onset and offset of each bird. All linear models were run using the function lm in the library *stats* in R.

To test for variation in the levels of the individual metabolites identified by the LC-MS, we used all data, including the replicated mid-day samples (total n = 64). Data from the subset of birds sampled both, two days before the culling and during the culling (n = 12; Table S3) were highly correlated ($r=0.92$ and $p<0.001$, Fig. S8). We ran independent LMMs for each metabolite, with metabolite levels as response variable (normalized), and treatment, time of day and their interaction as explanatory variables. ID was always included as random factor. We corrected the p-values of all these models by the false discovery rate test. Moreover, we ran two principal component analyses using the function prcomp in the library *stats* in R. For these, we used only the 73 metabolites found to be significantly affected by the treatment*time interaction. The two PCAs were run on the individual samples collected at mid-day or midnight, respectively. We then used the first two principal components (PC1 and PC2) of the midnight based PCA as explanatory variables in two linear models with onset and offset of activity as response variables, respectively.

### Supplementary figures

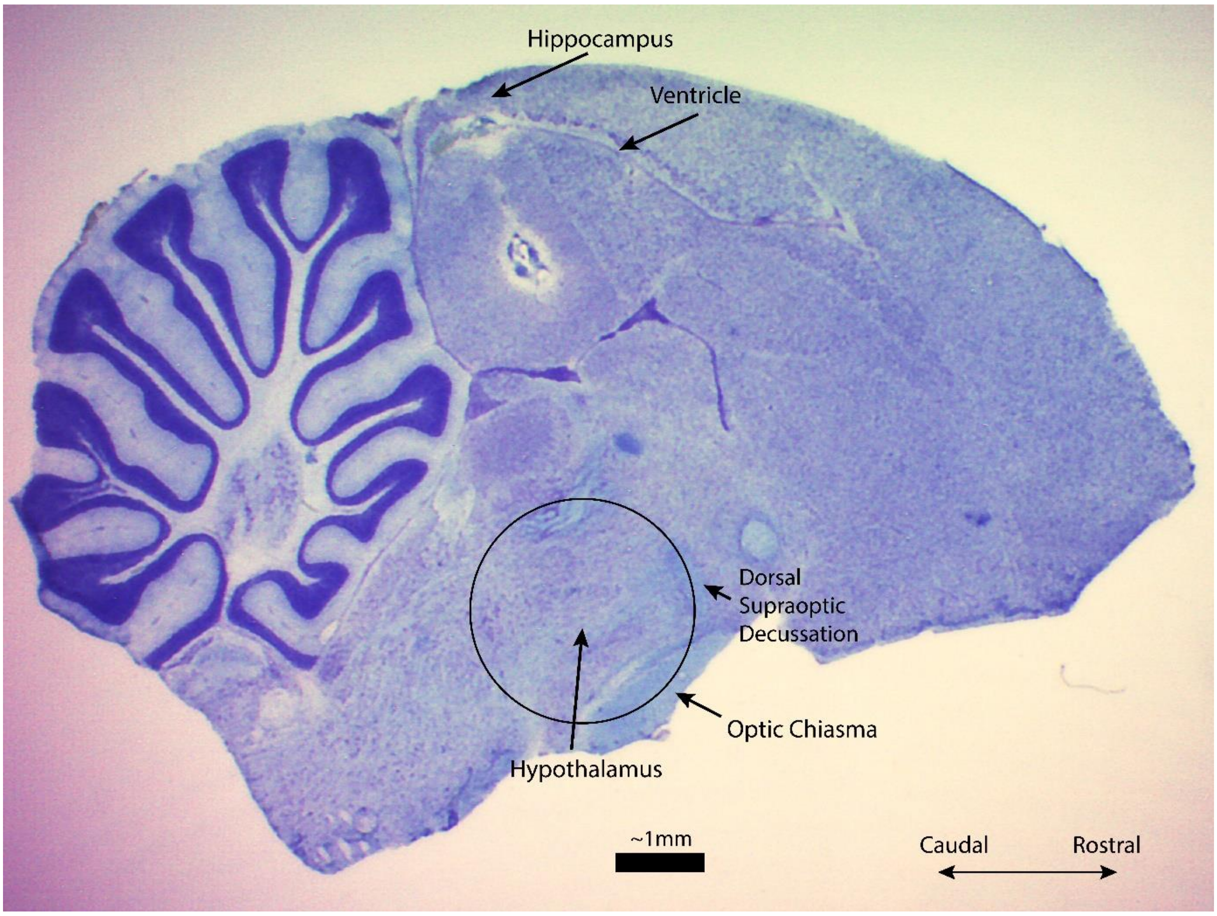

**Figure S1**. Sagittal section of a great tit brain sampled in the experiment. Highlighted are the two areas used in the analyses: the hypothalamus, sampled via a 3 mm punch in the region just dorso-caudal of the optic chiasm, caudal of the dorsal supraoptic decussation, and the hippocampus, the central superior region of the section delimited by the below lateral ventricle.

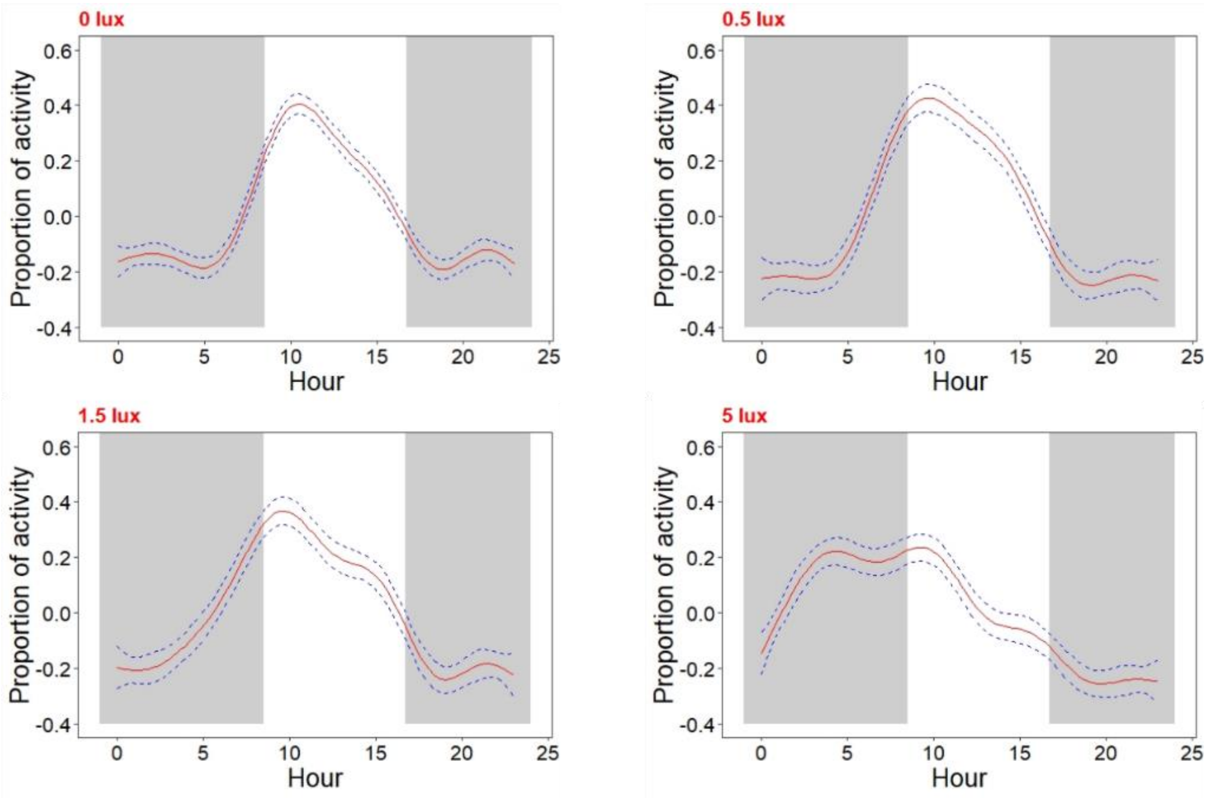

**Figure S2.** GAMM predictions for each light treatment for the proportion of active 2-min intervals per hour. In each panel, red line depicts predicted mean and blue dashed lines depict 95 % confidence intervals. Grey areas represent night hours, white areas represent daytime.

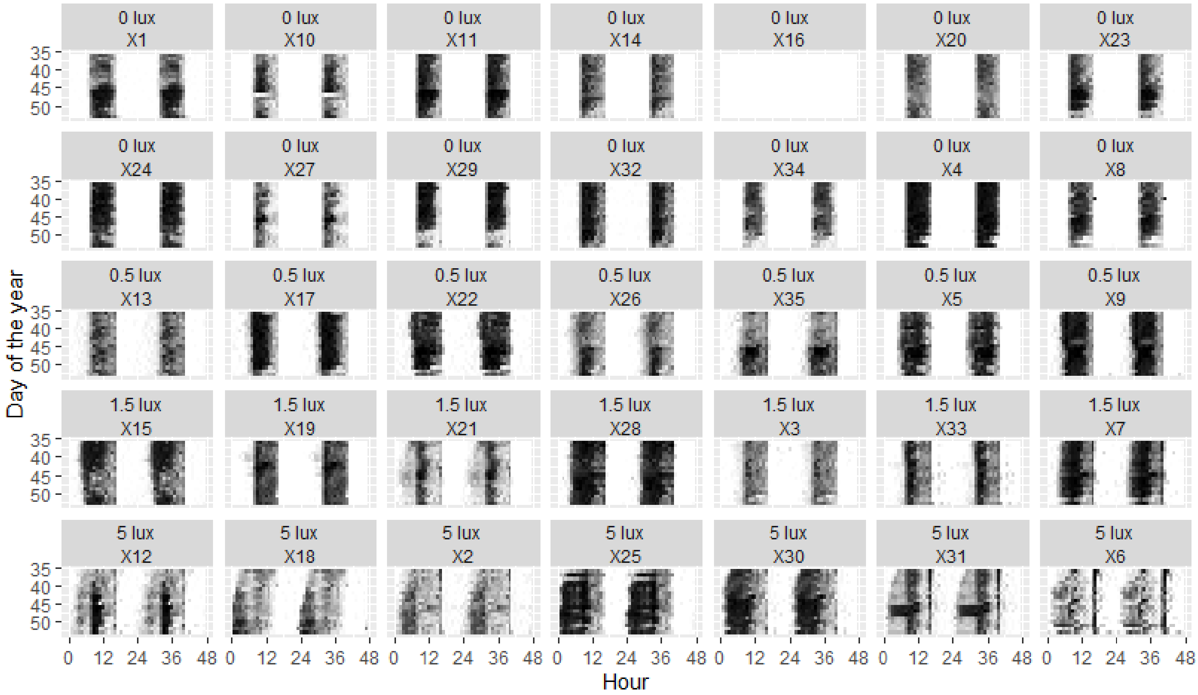

**Figure S3.** Actograms of all birds used in the experiment. The first two rows represent birds in the 0 lx treatment, the third row birds in the 0.5 lux treatment, the fourth row those in the 1.5 lux treatment, and the last row the birds in the 5 lux treatment. One bird (X16) died before the start of the experiment. Within each plot, rows represent days since the start of the year, and columns the hours of day. The intensity of black represents the amount of activity within each hour bin. Each actogram is double plotted to better visualize free-running rhythms of activity.

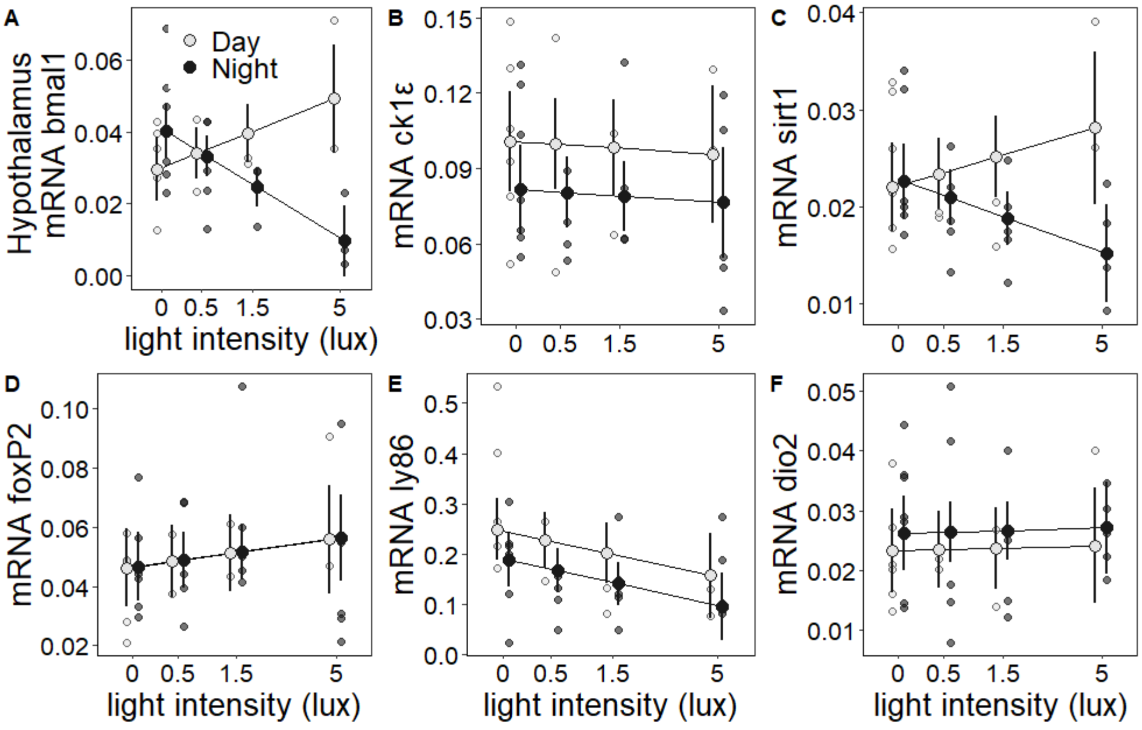

Figure S4. Changes in hypothalamic gene expression in response to ALAN of different intensity measured at mid-day vs. midnight. Large symbols ± SEM connected by lines represent model estimates, whereas small symbols depict raw data points (grey = mid-day, black = midnight).

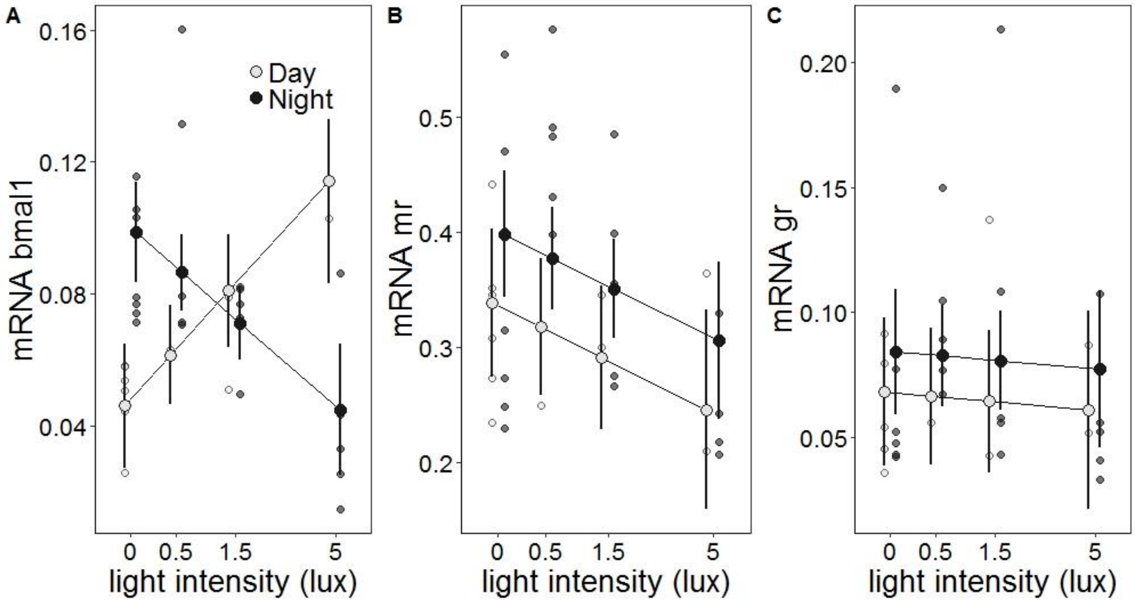

**Figure S5.** Changes in hippocampal gene expression in response to ALAN of different intensity. Large symbols ± SEM connected by lines represent model estimates, whereas small symbols depict raw data points (grey = mid-day, black = midnight).

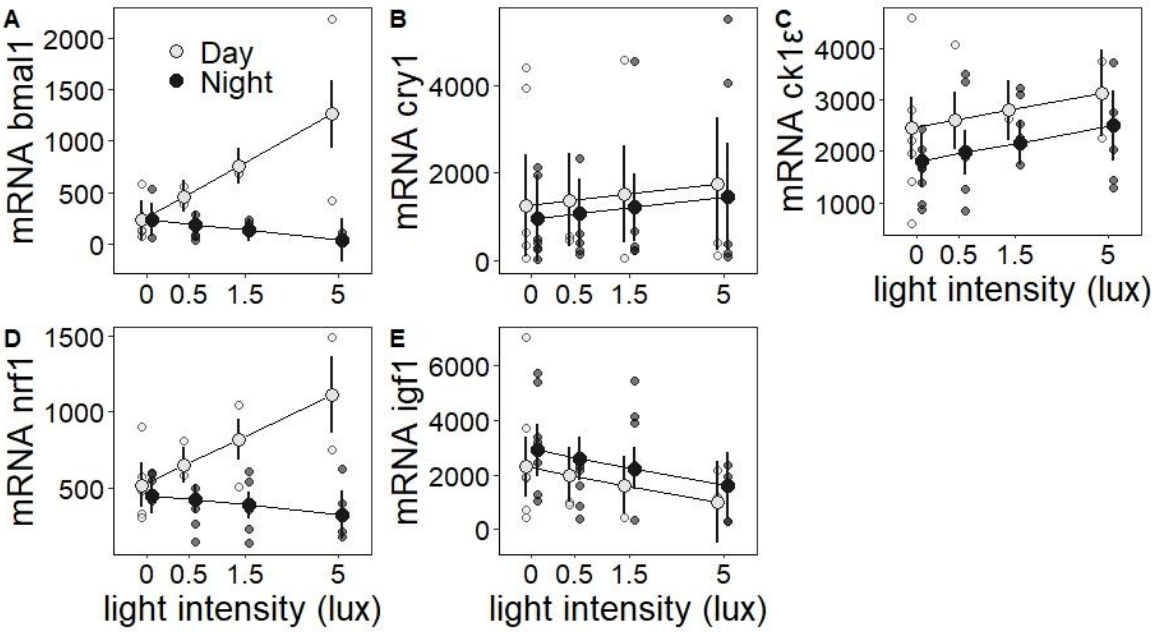

**Figure S6.** Changes in liver gene expression in response to ALAN of different intensity. One outlier was detected for *CRY1 (panel B)*, however, its exclusion did not qualitatively modify the statistical results. Large symbols ± SEM connected by lines represent model estimates, whereas small symbols depict raw data points (grey = mid-day, black = midnight).

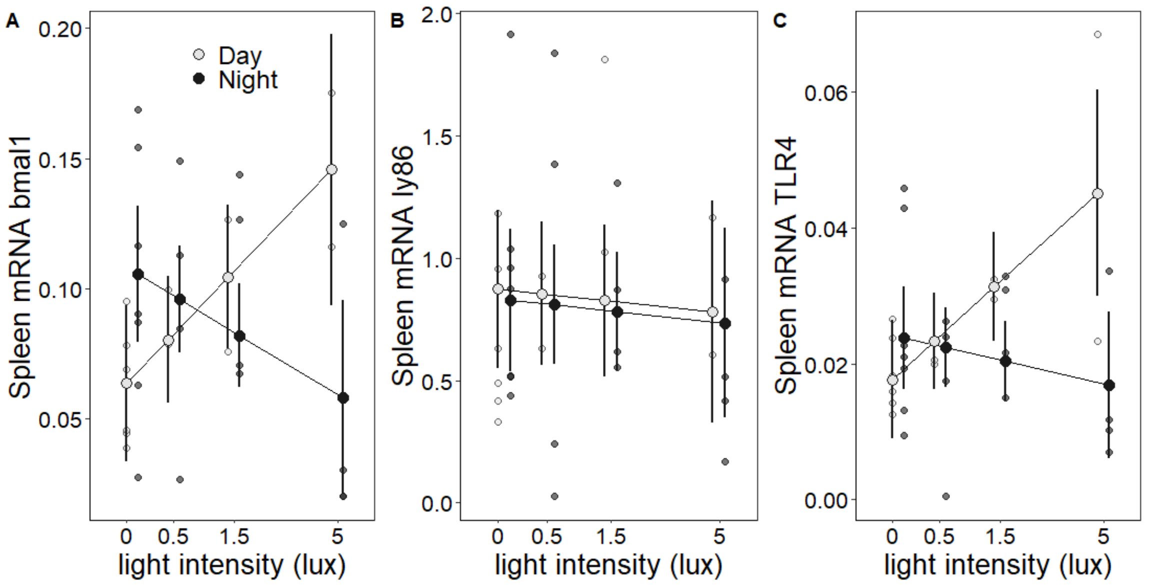

**Figure S7.** Changes in spleen gene expression in response to ALAN of different intensity. Large symbols ± SEM connected by lines represent model estimates, whereas small symbols depict raw data points (grey = mid-day, black = midnight).

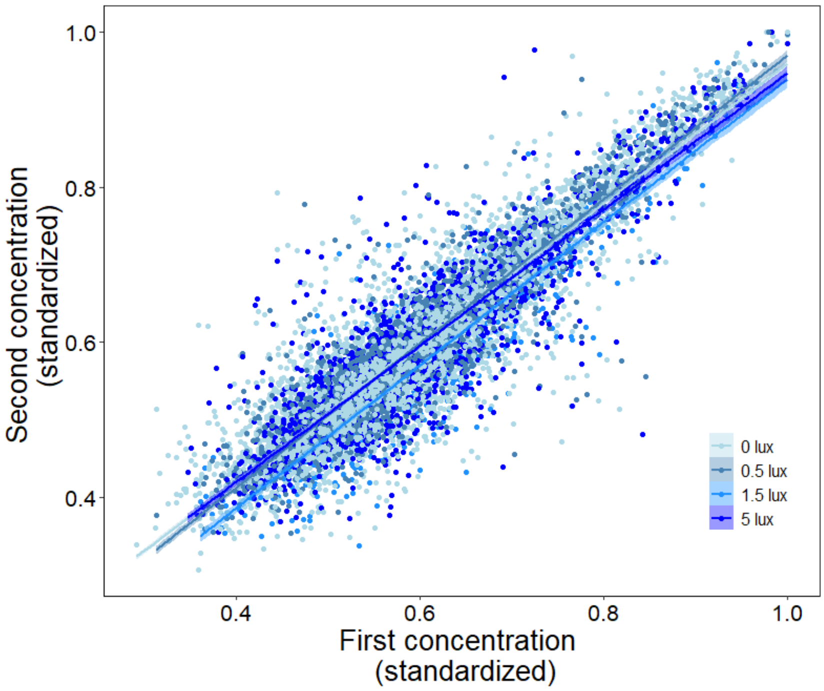

**Figure S8.** Within-individual correlations between standardised metabolite concentration considering the whole metabolome (892 metabolites). Each point represents one metabolite measured in one individual two days apart, before the final culling (x-axis) and at the time of culling (y-axis).

### Supplementary tables

**Table S1.** Overview of genes analyzed. Indicated are genes with their full names, a functional note, the tissues where they were measured (Hth=hypothalamus, Hip=hippocampus, Liv=liver, Spl=spleen), and forward and reverse primers for RT-qPCR. Superscripts indicate functional groups. [a] clock genes and modulators; [b] seasonal regulators; [c] metabolic genes; [d] immune genes; [e] endocrine genes; [f] reference genes.

| Gene | Full name | Functional note | Hth | Hip | Liv | Spl | Forward | Reverse |
|---|---|---|---|---|---|---|---|---|
| [a] *bmal1 (arntl)* | Brain and Muscle ARNT-Like1 | core clock gene | yes | yes | yes | yes | cgcttcgtggtgctacaaac | ccatctgctgccctgagaat |
| [a] *cry1* | Cryptochrome Circadian Regulator 1 | core clock gene | | | yes | | tcaccatttagcccggcatgc | gaaagaaggaactacaggacagccacatc |
| [a] *ck1ε (CSNK1E)* | Casein kinase I isoform epsilon | posttranslational modulator of clock proteins (PER) | yes | | yes | | cattaagtggtgtggagcag | aaatttgcgggaacagaagt |
| [b] *dio2* | Type II iodothyronine deiodinase | converts thyroxine (T4) to bioactive thyroid hormone triiodothyronine (T3) | yes | | | | tccacacttgccaccaacat | caaactgggaggagaagccc |
| [b] *foxP2* | Forkhead box protein P2 | transcription factor involved in vocal learning and plasticity | yes | | | | aaaggagcagtatggacagt | agctggtgggtatgtttttc |
| [c] *sirt1* | Sirtuin 1 | enzyme deacetylating transcription factors (clock-, stress-, ageing- linked) | yes | | | | gtcacaagttcatcgctttg | atcctttggattcctgcaac |
| [c] *nrf1* | Nuclear Respiratory Factor 1 | key nuclear transcription factor of involved in mitochondrial activity | | | yes | | gggccacgctggatgagtaca | gccagcgccgattccagat |
| [d] *ly86 (MD1)* | Lymphocyte antigen 86 | regulates T cell activation and cytokine production | yes | | | yes | gaccattgtgctgatattgcaaccc | agcttatcatgacccggccca |
| [d] *tlr4* | Toll-like receptor 4 | Pattern recognition factor, innate immune system activation | | | | yes | cacctccacaccttggatatt | tcgaaggtcaggagcttattg |
| [e] *nr3c1 (gr)* | Nuclear receptor subfamily 3, group C, member 1 | glucocorticoid receptor | | yes | | | attggctccgctgggaacg | aggcctcgtcagagcacacca |
| [e] *nr3c2 (mr)* | Nuclear receptor subfamily 3, group C, member 2 | mineralocorticoid receptor | | yes | | | tgtgtctgtcatcgtttgccttgag | cggacgaactgcaggctgatct |
| [e] *igf1* | Insulin-like growth factor 1 | hormone linked to mitochondrial biogenesis, respiration and ageing | | | yes | | ttgctgctggcccagaaacac | cacaactctggaagcagcattcatcc |
| [f] *RPL13* | ribosomal protein L13 | component of ribosome | yes | yes | yes | | tactccttcagcctctgcac | acaagaagtttgcccggact |
| [f] *RPL19* | ribosomal protein L19 | component of ribosome | yes | | | yes | ctgcggcaagaagaaggtgt | tcagcccatccttgatcagc |
| [f] *SDHA* | Succinate Dehydrogenase Complex Flavoprotein Subunit | mitochondrial respiratory chain | | yes | yes | yes | gggcaataactccacggcat | ttgtatggcaggtctctacga |
| [f] *PMM1* | Phosphomannomutase | glycosylation | | | yes | | caccccagaggagcgaatcga | tcgagcacattgaggcagtagcg |
| [f] *TBP* | TATA-binding protein | general transcription factor | | | yes | | aaaactattgcacttcgtgcccga | gaatatcagtgcagtggtacgtggttctct |

**Table S2.** Sample sizes for treatment and time of sampling used in the gene expression analyses.

| Treatment | Day | Night |
|---|---|---|
| 0 lux | 6 | 7 |
| 0.5 lux | 2 | 5 |
| 1.5 lux | 2 | 5 |
| 5 lux | 2 | 5 |

**Table S3.** Sample sizes for treatment and time of sampling used for the metabolomics analysis. All birds were sampled at mid-day two days before the final culling, on Feb 20$^{th}$. Birds were then re-sampled at mid-day or midnight during the culling. Metabolomics data from 64 individual bird samples (4 samples were excluded due to low signal in the LC-MS data) were used in our analyses. For a subset of birds (n = 12, the sum of all birds sampled at mid-day during culling), we have two mid-day samples which we used to examine within-individual correlation of metabolite levels (see Fig. S8 for these results).

| Treatment | Pre-culling samples (Feb 20$^{th}$ 2014) | Culling samples (Feb 22$^{nd}$ & 23$^{rd}$ 2014) | |
|---|---|---|---|
| | Day | Day | Night |
| 0 lux | 13 | 6 | 6 |
| 0.5 lux | 7 | 2 | 5 |
| 1.5 lux | 6 | 2 | 3 |
| 5 lux | 7 | 2 | 5 |

**Table S4.** Results of generalized additive mixed model (GAMM) testing for variation in the proportion of 2-min intervals spent active per hour of day, depending on night light intensity. Treatment was included as linear predictor. Hour of day was included as smoothed predictor, and we included four smoothed terms referring to the four treatments. Plots for these smoothed terms can be seen in Fig. 2 of the main text.

| **Parametric coefficients** | | | | |
|---|---|---|---|---|
| *Predictor* | *Estimate* | *Std. Error* | *t value* | *p value* |
| Intercept | 0.18 | 0.01 | 14.51 | < 0.001 |
| Treatment | 0.02 | 0.01 | 3.18 | **0.001** |
| **Significance of smooth terms:** | | | | |
| *Predictor* | *F* | *p value* | | |
| s(Hour of Day)*Treatment 0 lux | 177.38 | **<0.001** | | |
| s(Hour of Day)*Treatment 0.5 lux | 141.02 | **<0.001** | | |
| s(Hour of Day)*Treatment 1.5 lux | 95.86 | **<0.001** | | |
| s(Hour of Day)*Treatment 5 lux | 81.13 | **<0.001** | | |

**Table S5.** Results of Gaussian linear mixed models testing for variation in activity levels depending on night light intensity, day of the experiment and their interaction. The day of the experiment was also coded as a quadratic term for testing non-linear changes in activity traits over the course of the experiment after the initial exposure to ALAN. The error structure used in each model is specified in brackets.

| **Onset of activity** | | | | |
|---|---|---|---|---|
| *predictor* | *estimate* | *std.error* | *t value* | *p value* |
| Intercept | -134.7 | 10.4 | -12.92 | <0.001 |
| Day | -23.7 | 9.8 | -2.41 | **0.017** |
| Treatment | -124.9 | 10.4 | -11.96 | **<0.001** |
| $Day^2$ | 19 | 9.3 | 2.04 | **0.042** |
| Treatment*Day | -34.6 | 9.9 | -3.52 | **<0.001** |
| $Treatment*Day^2$ | 19.4 | 9.3 | 2.08 | **0.038** |
| **Offset of activity** | | | | |
| *predictor* | *estimate* | *std.error* | *t value* | *p value* |
| Intercept | 4.2 | 7.5 | 0.56 | 0.578 |
| Day | -8 | 7.6 | -1.06 | 0.289 |
| Treatment | 8.4 | 7.5 | 1.12 | 0.271 |
| $Day^2$ | 4.5 | 7.6 | 0.59 | 0.556 |
| Treatment*Day | -10.9 | 1.8 | -6.15 | **<0.001** |
| **Nocturnal activity** | | | | |
| *predictor* | *estimate* | *std.error* | *t value* | *p value* |
| Intercept | -1.7 | 0.1 | -12.1 | **<0.001** |
| Day | 0.1 | 0.1 | 0.51 | **<0.001** |
| Treatment | 1.1 | 0.1 | 8.83 | **<0.001** |
| **Total 24h activity** | | | | |
| *predictor* | *estimate* | *std.error* | *t value* | *p value* |
| Intercept | 377.7 | 17.3 | 21.78 | **<0.001** |
| Day | 41.9 | 10.8 | 3.87 | **<0.001** |
| Treatment | 32.8 | 17.4 | 1.89 | 0.068 |
| $Day^2$ | -30.9 | 10.8 | -2.86 | **0.004** |
| Treatment*Day | 14.6 | 2.5 | 5.77 | **<0.001** |

**Table S6.** Estimated period length (tau) of Great tits exposed to ALAN until activity patterns stabilized. Shown are estimated means from a Gaussian LM and outcomes of Tukey post-hoc testing for differences in tau through pair-wise contrasts of treatment groups. Tau was estimated via Lomb-Scargle periodogram analysis implemented in the software Chronoshop (courtesy of Kamiel Spoelstra), using only the first 10 days of activity data. During this phase shifting interval, period lengths in the 5 lx group were similar to the reported free-running period length of this species [13].

| **Estimated means** | | | |
|---|---|---|---|
| *treatment* | *estimated mean (mins)* | *lower 95 % CI* | *upper 95% CI* |
| 0 lux | 23.95 | 23.88 | 24.01 |
| 0.5 lux | 23.89 | 23.80 | 23.97 |
| 1.5 lux | 23.85 | 23.76 | 23.94 |
| 5 lux | 23.59 | 23.50 | 23.68 |
| **Post-hoc test** | | | |
| *contrast* | *estimated difference (mins)* | *std.error* | *P value* |
| 0 lux - 0.5 lux | 3.8 | 3.2 | 0.646 |
| 0 lux - 1.5 lux | 6.1 | 3.2 | 0.254 |
| 0 lux - 5 lux | 21.5 | 3.2 | **<0.001** |
| 0.5 lux - 1.5 lux | 2.3 | 3.7 | 0.924 |
| 0.5 lux - 5 lux | 17.7 | 3.7 | **<0.001** |
| 1.5 lux - 5 lux | 15.4 | 3.7 | **0.001** |

**Table S7.** Results of linear models (Gaussian error structure) testing for variation in mRNA levels of six different genes in the hypothalamus. Estimates for the predictor sampling time refer to midnight values, while mid-day values are the reference level.

**HYPOTHALAMUS GENE EXPRESSION**

**bmal1**

| *predictor* | *estimate* | *std.error* | *t value* | *p value* |
|---|---|---|---|---|
| Intercept | 0.03 | 0.004 | 6.8 | <0.001 |
| Treatment | 0.011 | 0.005 | 2.1 | **0.044** |
| Time | 0.01 | 0.006 | 1.84 | 0.076 |
| Treatment*Time | -0.028 | 0.006 | -4.37 | **<0.001** |

**ck1ε**

| *predictor* | *estimate* | *std.error* | *t value* | *p value* |
|---|---|---|---|---|
| Intercept | 0.101 | 0.01 | 10.32 | **<0.001** |
| Treatment | -0.003 | 0.008 | -0.38 | 0.709 |
| Time | -0.019 | 0.011 | -1.72 | 0.095 |

**sirt1**

| *predictor* | *estimate* | *std.error* | *t value* | *p value* |
|---|---|---|---|---|
| Intercept | 0.022 | 0.002 | 9.83 | <0.001 |
| Treatment | 0.003 | 0.003 | 1.26 | 0.216 |
| Time | 0.001 | 0.003 | 0.22 | 0.831 |
| Treatment*Time | -0.008 | 0.003 | -2.29 | **0.029** |

**dio2**

| *predictor* | *estimate* | *std.error* | *t value* | *p value* |
|---|---|---|---|---|
| Intercept | 0.023 | 0.003 | 6.75 | <0.001 |
| Treatment | 0 | 0.003 | 0.17 | 0.866 |
| Time | 0.003 | 0.004 | 0.76 | 0.455 |

**foxp2**

| *predictor* | *estimate* | *std.error* | *t value* | *p value* |
|---|---|---|---|---|
| Intercept | 0.046 | 0.007 | 7.07 | <0.001 |
| Treatment | 0.005 | 0.005 | 1.02 | 0.317 |
| Time | 0 | 0.007 | 0.05 | 0.958 |

**ly86**

| *predictor* | *estimate* | *std.error* | *t value* | *p value* |
|---|---|---|---|---|
| Intercept | 0.249 | 0.03 | 8.29 | <0.001 |
| Treatment | -0.051 | 0.024 | -2.12 | **0.042** |
| Time | -0.061 | 0.034 | -1.79 | 0.083 |

**Table S8.** Results of linear models (Gaussian error structure) testing for variation in mRNA levels of three different genes in the hippocampus. Estimates for the predictor sampling time refer to midnight values, while mid-day values are the reference level.

| **HIPPOCAMPUS GENE EXPRESSION** | | | | |
|---|---|---|---|---|
| **bmal1** | | | | |
| *predictor* | *estimate* | *std.error* | *t value* | *p value* |
| Intercept | 0.046 | 0.009 | 5.03 | <0.001 |
| Treatment | 0.038 | 0.011 | 3.59 | **0.001** |
| Time | 0.053 | 0.012 | 4.45 | **<0.001** |
| Treatment*Time | -0.068 | 0.013 | -5.22 | **<0.001** |
| **mineralocorticoid receptor** | | | | |
| *predictor* | *estimate* | *std.error* | *t value* | *p value* |
| Intercept | 0.338 | 0.032 | 10.69 | <0.001 |
| Treatment | -0.052 | 0.025 | -2.11 | **0.044** |
| Time | 0.06 | 0.036 | 1.67 | 0.105 |
| **glucocorticoid receptor** | | | | |
| *predictor* | *estimate* | *std.error* | *t value* | *p value* |
| Intercept | 0.068 | 0.015 | 4.67 | <0.001 |
| Treatment | -0.004 | 0.011 | -0.35 | 0.728 |
| Time | 0.016 | 0.016 | 0.99 | 0.329 |

**Table S9.** Relationships between *BMAL1* mRNA levels in different tissues. Shown are results of Gaussian linear models testing for the relationship between mRNA levels (all log-transformed) in two tissues per model, while controlling for sampling time and treatment, which were included as covariates in all models.

| ***bmal1* hippocampus ~ *bmal1* hypothalamus** | | | | |
|---|---|---|---|---|
| *Predictor* | *Estimate* | *Std. Error* | *t value* | *p value* |
| Intercept | -0.44 | 0.30 | -1.50 | 0.145 |
| bmal1 hypothalamus | 0.71 | 0.09 | 8.02 | **< 0.001** |
| Time | 0.40 | 0.11 | 3.72 | 0.001 |
| Treatment | 0.04 | 0.03 | 1.44 | 0.160 |
| ***bmal1* liver ~ *bmal1* hypothalamus** | | | | |
| *Predictor* | *Estimate* | *Std. Error* | *t value* | *p value* |
| Intercept | 9.62 | 0.77 | 12.57 | < 0.001 |
| bmal1 hypothalamus | 1.11 | 0.23 | 4.89 | **< 0.001** |
| Time | -0.80 | 0.27 | -2.97 | 0.006 |
| Treatment | 0.04 | 0.08 | 0.58 | 0.567 |
| ***bmal1* spleen ~ *bmal1* hypothalamus** | | | | |
| *Predictor* | *Estimate* | *Std. Error* | *t value* | *p value* |
| Intercept | -2.17 | 0.43 | -5.10 | < 0.001 |
| bmal1 hypothalamus | 0.43 | 0.16 | 2.72 | **0.011** |
| Time | -0.28 | 0.20 | -1.41 | 0.170 |
| Treatment | -0.13 | 0.05 | -2.46 | 0.021 |
| ***bmal1* spleen ~ *bmal1* liver** | | | | |
| *Predictor* | *Estimate* | *Std. Error* | *t value* | *p value* |
| Intercept | -4.71 | 0.65 | -7.30 | < 0.001 |
| bmal1 liver | 0.37 | 0.10 | 3.59 | **0.001** |
| Time | 0.32 | 0.23 | 1.42 | 0.166 |
| Treatment | 0.00 | 0.05 | 0.02 | 0.987 |

**Table S10.** Results of linear models (Gaussian error structure) testing for variation in mRNA levels of four different genes in the liver. Estimates for the predictor sampling time refer to midnight values, while mid-day values are the reference level.

| **LIVER GENE EXPRESSION** | | | | |
|---|---|---|---|---|
| **bmal1** | | | | |
| *predictor* | *estimate* | *std.error* | *t value* | *p value* |
| Intercept | 228.703 | 94.065 | 2.43 | 0.021 |
| Treatment | 576.357 | 112.237 | 5.14 | **<0.001** |
| Time | 4.893 | 122.818 | 0.04 | 0.968 |
| Treatment*Time | -684.509 | 138.231 | -4.95 | **<0.001** |
| **ck1ε** | | | | |
| *predictor* | *estimate* | *std.error* | *t value* | *p value* |
| Intercept | 2446.672 | 300.406 | 8.14 | <0.001 |
| Treatment | 376.241 | 242.042 | 1.55 | 0.131 |
| Time | -625.039 | 342.306 | -1.83 | 0.078 |
| **nrf1** | | | | |
| *predictor* | *estimate* | *std.error* | *t value* | *p value* |
| Intercept | 516.546 | 71.402 | 7.23 | <0.001 |
| Treatment | 330.961 | 85.196 | 3.88 | **0.001** |
| Time | -65.376 | 93.228 | -0.7 | 0.489 |
| Treatment*Time | -402.217 | 104.928 | -3.83 | **0.001** |
| **igf1** | | | | |
| *predictor* | *estimate* | *std.error* | *t value* | *p value* |
| Intercept | 2289.529 | 541.793 | 4.23 | <0.001 |
| Treatment | -720.868 | 436.532 | -1.65 | 0.109 |
| Time | 604.69 | 617.361 | 0.98 | 0.335 |

**Table S11.** Results of linear models (Gaussian error structure) testing for variation in mRNA levels of three different genes in the spleen. Estimates for the predictor sampling time refer to midnight values, while mid-day values are the reference level.

| SPLEEN GENE EXPRESSION | | | | |
|---|---|---|---|---|
| **bmal1** | | | | |
| *predictor* | *estimate* | *std.error* | *t value* | *p value* |
| Intercept | 0.063 | 0.015 | 4.3 | 0 |
| Treatment | 0.046 | 0.018 | 2.61 | **0.015** |
| Time | 0.043 | 0.019 | 2.19 | **0.037** |
| Treatment*Time | -0.072 | 0.022 | -3.25 | **0.003** |
| **ly86** | | | | |
| *predictor* | *estimate* | *std.error* | *t value* | *p value* |
| Intercept | 0.873 | 0.158 | 5.52 | 0 |
| Treatment | -0.049 | 0.132 | -0.37 | 0.711 |
| Time | -0.045 | 0.183 | -0.25 | 0.806 |
| **tlr4** | | | | |
| *predictor* | *estimate* | *std.error* | *t value* | *p value* |
| Intercept | 0.018 | 0.004 | 4.09 | 0 |
| Treatment | 0.015 | 0.005 | 2.98 | **0.006** |
| Time | 0.006 | 0.006 | 1.12 | 0.272 |
| Treatment*Time | -0.019 | 0.006 | -2.97 | **0.006** |

**Table S12.** List of 29 metabolites significantly affected by the linear effect of treatment, as tested via individuals LMMs run on all 755 identified metabolites. P-values were corrected using a false discovery rate of 0.05.

| metabolite | F value | p value | p (fdr) |
|---|---|---|---|
| β-ketophosphonate | 12 | 0.002 | 0.013 |
| [FA] O-Palmitoyl-R-carnitine | 7.7 | 0.007 | 0.035 |
| [PC (20:0)] 1-eicosanoyl-sn-glycero-3-phosphocholine | 11.6 | 0.002 | 0.012 |
| [PC acetyl(17:2)] 1-heptadecyl-2-acetyl-sn-glycero-3-phosphocholine | 10.4 | 0.003 | 0.017 |
| 1,2-dioctanoyl-1-amino-2,3-propanediol | 11 | 0.002 | 0.014 |
| 3-4-DihydroxyphenylglycolO-sulfate | 7.4 | 0.009 | 0.039 |
| 4-Pyridoxate | 8.7 | 0.006 | 0.03 |
| Ala-Asp-Pro | 8.4 | 0.007 | 0.033 |
| Anandamide | 12.8 | 0.001 | 0.005 |
| D-Glucuronate | 12.8 | 0.001 | 0.008 |
| D-Ornithine | 7.4 | 0.008 | 0.038 |
| dimethylsulfonio-2-hydroxybutyrate | 7.7 | 0.01 | 0.042 |
| gamma-L-Glutamylputrescine | 8.7 | 0.004 | 0.024 |
| Glu-Arg | 9.4 | 0.003 | 0.018 |
| Glu-Thr | 7.7 | 0.007 | 0.034 |
| Glyceraldehyde | 7.1 | 0.01 | 0.042 |
| Hypotaurine | 11.7 | 0.002 | 0.011 |
| Imidazole-4-acetate | 10.9 | 0.002 | 0.01 |
| L-a-glutamyl-L-Lysine | 9.6 | 0.003 | 0.017 |
| L-Aspartate | 7.3 | 0.009 | 0.04 |
| L-Cysteinylglycinedisulfide | 6.8 | 0.009 | 0.041 |
| Leucyl-leucine | 6.9 | 0.011 | 0.046 |
| LysoPE(0:0/22:0) | 8.8 | 0.006 | 0.028 |
| Ne,Ne dimethyllysine | 7.3 | 0.009 | 0.039 |
| Protoporphyrin | 7.1 | 0.01 | 0.042 |
| Stearoylcarnitine | 9.3 | 0.003 | 0.019 |
| Taxa-4(20),11(12)-dien-5alpha-yl acetate | 8.9 | 0.004 | 0.022 |
| Tetradecanoylcarnitine | 12.2 | 0.001 | 0.01 |
| Xylitol | 6.9 | 0.011 | 0.046 |

**Table S13.** List of 73 metabolites significantly affected by the interaction of treatment and sampling time, as tested via individuals LMMs run on all 755 identified metabolites. P-values were corrected using a false discovery rate of 0.05.

| metabolite | F value | p value | p (fdr) |
|---|---|---|---|
| (S)-ATPA | 7.9 | 0.007 | 0.027 |
| [FA (12:3)] 3,6,8-dodecatrien-1-ol | 4.5 | 0.039 | 0.047 |
| [FA (14:2)] 5,8-tetradecadienoic acid | 4.5 | 0.04 | 0.047 |
| [FA (16:2)] 9,12-hexadecadienoic acid | 5.4 | 0.025 | 0.044 |
| [FA (18:1)] 9Z-octadecenoic acid | 6.2 | 0.017 | 0.041 |
| [FA (20:0)] 11Z-eicosenoic acid | 9.8 | 0.003 | 0.026 |
| [FA (20:0)] eicosanoic acid | 4.6 | 0.037 | 0.047 |
| [FA (23:0/2:0)] Tricosanedioic acid | 8.1 | 0.007 | 0.027 |
| [FA methyl(18:0)] 11R,12S-methylene-octadecanoic acid | 5.8 | 0.021 | 0.042 |
| [FA methyl,oxo(5:0/2:0)] 2-methylene-4-oxo-pentanedioic acid | 5 | 0.029 | 0.046 |
| [FA oxo(5:2/5:0/4:0)] (1S,2S)-3-oxo-2-pentyl-cyclopentanebutanoic acid | 4.8 | 0.034 | 0.047 |
| [FA oxo(5:2/5:0/6:0)] (1R,2R)-3-oxo-2-pentyl-cyclopentanehexanoic acid | 8.2 | 0.007 | 0.027 |
| [Fv (2:0)] Flavaprenin 7,4'-diglucoside | 10.4 | 0.002 | 0.024 |
| [PE (16:0)] 1-hexadecanoyl-sn-glycero-3-phosphoethanolamine | 4.6 | 0.037 | 0.047 |
| [PE (17:1)] 1-(9Z-heptadecenoyl)-sn-glycero-3-phosphoethanolamine | 4.2 | 0.044 | 0.047 |
| [SP] Sphinganine-1-phosphate | 7.6 | 0.008 | 0.03 |
| 2-Aminomuconate | 9.1 | 0.005 | 0.027 |
| 2-Hydroxyethanesulfonate | 5.4 | 0.024 | 0.043 |
| 2-monooleoylglycerol | 5.9 | 0.019 | 0.042 |
| 2-thiouridine | 6.6 | 0.012 | 0.038 |
| 2,7-Anhydro-alpha-N-acetylneuraminic acid | 6.6 | 0.013 | 0.039 |
| 3-4-DihydroxyphenylglycolO-sulfate | 4.7 | 0.036 | 0.047 |
| 3-Aminopropanesulfonate | 11.5 | 0.001 | 0.022 |
| 3-Dehydroxycarnitine | 6.5 | 0.015 | 0.041 |
| 3-Methylguanine | 4.3 | 0.045 | 0.047 |
| 3-Oxododecanoic acid | 5.1 | 0.029 | 0.046 |
| 3-sulfopropanoate | 4.1 | 0.048 | 0.048 |
| 5-Amino-4-chloro-2-(2,3-dihydroxyphenyl)-3(2H)-pyridazinone | 5.5 | 0.022 | 0.043 |
| 9-Decenoylcarnitine | 8.2 | 0.007 | 0.027 |
| Ala-Ser | 6.2 | 0.016 | 0.041 |
| D-Glucarate | 4.3 | 0.043 | 0.047 |
| D-Proline | 4.7 | 0.037 | 0.047 |
| Ethanolamine phosphate | 8 | 0.006 | 0.027 |
| Gabapentin | 5.7 | 0.021 | 0.042 |
| gamma-L-Glutamylputrescine | 13 | 0.001 | 0.022 |
| Glu-Arg | 5.5 | 0.023 | 0.043 |
| Glu-Pro | 11.3 | 0.002 | 0.022 |
| Glu-Thr | 4.1 | 0.049 | 0.049 |
| Glutathione disulfide | 4.1 | 0.047 | 0.048 |
| Glycylproline | 4.5 | 0.04 | 0.047 |
| Homoarginine | 4.6 | 0.038 | 0.047 |
| Homocysteinesulfinicacid | 4.7 | 0.034 | 0.047 |

| | | | |
|---|---|---|---|
| hydrogen iodide | 4.5 | 0.038 | 0.047 |
| Imidazole-4-acetate | 4.6 | 0.035 | 0.047 |
| L-a-glutamyl-L-Lysine | 6 | 0.017 | 0.041 |
| L-Arginine | 5.4 | 0.025 | 0.044 |
| L-Citrulline | 4.5 | 0.04 | 0.047 |
| L-Glutamate | 5.1 | 0.027 | 0.045 |
| L-Lysine | 7 | 0.012 | 0.038 |
| L-Threonine | 5.7 | 0.019 | 0.042 |
| L-Tyrosine | 12.6 | 0.001 | 0.022 |
| Leu-Asn-His | 6.3 | 0.016 | 0.041 |
| Leu-Phe-Cys | 4.3 | 0.043 | 0.047 |
| Linoelaidylcarnitine | 4.1 | 0.047 | 0.048 |
| Linoleate | 8.9 | 0.005 | 0.027 |
| LysoPC(22:4(7Z,10Z,13Z,16Z)) | 4.6 | 0.038 | 0.047 |
| LysoPC(22:5(4Z,7Z,10Z,13Z,16Z)) | 10.7 | 0.002 | 0.022 |
| LysoPE(0:0/22:5(4Z,7Z,10Z,13Z,16Z)) | 4.3 | 0.042 | 0.047 |
| N'-Phosphoguanidinoethyl methyl phosphate | 7.9 | 0.007 | 0.027 |
| N-Acetyl-D-fucosamine | 9.1 | 0.005 | 0.027 |
| N-Acetyl-L-aspartate | 6.2 | 0.015 | 0.041 |
| N-Acetylneuraminate | 7.1 | 0.01 | 0.034 |
| N6-Methyl-L-lysine | 4.6 | 0.038 | 0.047 |
| omega-Cyclohexylundecanoic acid | 6.1 | 0.018 | 0.042 |
| Phenylacetic acid | 5.2 | 0.027 | 0.045 |
| Propanoyl phosphate | 4.2 | 0.044 | 0.047 |
| Quinalphos | 8 | 0.006 | 0.027 |
| Retronecine | 5.6 | 0.024 | 0.043 |
| S-Acetyldihydrolipoamide | 10.7 | 0.002 | 0.022 |
| Stachydrine | 10.2 | 0.003 | 0.026 |
| Sulfite | 5.7 | 0.02 | 0.042 |
| Tetradecanoylcarnitine | 5.1 | 0.029 | 0.046 |
| Thr-Asp-Pro | 4.3 | 0.043 | 0.047 |
